# Vagus nerve stimulation recruits the central cholinergic system to enhance perceptual learning

**DOI:** 10.1101/2022.01.28.478197

**Authors:** Kathleen A. Martin, Eleni S. Papadoyannis, Jennifer K. Schiavo, Saba Shokat Fadaei, Sofia Orrey Valencia, Nesibe Z. Temiz, Matthew J. McGinley, David A. McCormick, Robert C. Froemke

**Affiliations:** Skirball Institute for Biomolecular Medicine, New York University School of Medicine, New York, NY, USA; Neuroscience Institute, New York University School of Medicine, New York, NY, USA; Department of Otolaryngology, New York University School of Medicine, New York, NY, USA; Department of Neuroscience and Physiology, New York University School of Medicine, New York, NY, USA; Center for Neural Science, New York University, New York, NY, USA; Princeton Neuroscience Institute, Princeton University, Princeton, New Jersey; Friedrich Miescher Institute for Biomedical Research, Basel, Switzerland; Duncan Neurological Research Institute, Baylor College of Medicine, Houston TX 77030; Department of Neuroscience, Baylor College of Medicine, Houston TX 77030; Institute of Neuroscience, University of Oregon, Eugene OR 97403

## Abstract

Perception can be refined by experience up to certain limits. It is unclear if perceptual limitations are absolute or could be partially overcome via enhanced neuromodulation and/or plasticity. Recent studies suggest the utility of peripheral nerve stimulation - specifically vagus nerve stimulation (VNS) - for altering neural activity and augmenting experience-dependent plasticity, although little is known about central mechanisms recruited by VNS. Here we developed an auditory discrimination task for mice implanted with a VNS electrode. VNS occurring during behavior gradually improved discrimination abilities beyond the level achieved by training alone. Using two-photon imaging, we identified changes to auditory cortical responses and activation of cortically-projecting cholinergic axons with VNS. Anatomical and optogenetic experiments indicated that VNS could enhance task performance via activation of the central cholinergic system. These results highlight the importance of cholinergic modulation for the efficacy of VNS, perhaps enabling further refinement of VNS methodology for clinical conditions.

## Introduction

Sensory processing is refined over development and must continue to be adaptive throughout life, in order to adequately regulate behavior in dynamic and challenging environments. This requires that adult perceptual and cognitive abilities are not fixed, but rather can be improved with additional experience and training. Perceptual learning, i.e., improvement in sensory perceptual abilities with practice, has been shown to occur across a wide range of domains, including visual processing of orientation, binocularity, and other spatial features^1–7^, auditory processing of pitch and temporal intervals^8–12^, and the detection and recognition of sensory stimuli in other modalities^13–15^. However, changes in perceptual abilities can often be quite limited and stimulus-specific, even after extensive periods of training^5,16–18^. It is generally believed that adult perceptual learning across species relies on neural mechanisms of synaptic plasticity within the cerebral cortex^1,7,14,17^, but it remains unclear what factors might limit cortical plasticity achieved by perceptual training and any consequent changes in perception and behavior. Conversely, mechanisms for plasticity not engaged during training might then be available for recruitment by other means, and perhaps improve sensory perceptual abilities even further.

Studies of the mammalian auditory system have proved to be especially revealing for determining the mechanisms connecting perceptual learning and neural plasticity. A long literature has connected changes in auditory experience with long-lasting neural changes in the central auditory system. In particular, many studies have demonstrated how auditory training affects tonotopic maps and single neuron responses, showing that acoustic stimuli predictive of outcome generally have enhanced representations in the auditory cortex^8,9,11,12,19,20^. Despite clear evidence that behavioral training eventually affects cortical maps and receptive fields, it is less clear how these changes are initially induced and occur throughout experience and learning to produce perceptual changes^21,22^. One consistent finding across systems and species is that central neuromodulatory systems such as the cholinergic basal forebrain are important for sensory perceptual learning and cortical plasticity^23–38^. However, it remains unclear the extent to which modulatory systems are endogenously recruited under different contexts or forms of training, and if it might be possible to artificially leverage central modulation to further improve perceptual learning.

Peripheral nerve stimulation provides an opportunity to augment performance and lead to lasting improvements in auditory perceptual abilities via activation of neuromodulatory systems. Vagus nerve stimulation (VNS) has been used for several clinical applications in humans, including treating epilepsy^39,40^, alleviating treatment-resistant depression^41^, and motor deficits after stroke ^42^. Despite its well-established benefits, less is known about the circuit mechanism through which VNS acts. This has led to experimental approaches for studying VNS in a range of species including rats^43,44^, ferrets^45^, and non-human primates^46^. Recently, in collaboration we developed and validated a VNS cuff electrode for mice^47–50^ to investigate the role of neuromodulation in VNS-mediated motor and perceptual learning. Previous work in non-human model organisms has shown that vagus nerve stimulation can induce neuroplasticity in primary sensory areas^43,44,51^ and in motor areas^50,52,53^. This enhanced neuroplasticity is thought to be partially mediated by indirectly activating neuromodulatory networks, including noradrenergic neurons in locus coeruleus^49,54,55^ and cholinergic neurons in basal forebrain^48–50^ via the nucleus tractus solitarii (NTS) ^56,57^. With the development of VNS cuffs for mice, we aimed to study the circuit mechanisms and modulatory systems activated by VNS, asking how VNS might be applied to promote plasticity and improve auditory perceptual learning.

## Results

### Auditory perceptual training in head-fixed mice

We first developed an auditory task to parametrically quantify the psychophysical abilities of mice for frequency discrimination. A total of 38 mice (5 female wild-type, 11 male wild-type, 8 female ChAT-Cre, 14 male ChAT-Cre) were used for behavioral studies. Animals were head-fixed and progressively trained to classify presented pure tones in a two-alternative forced-choice (2AFC) task for a water reward. On each trial, mice were presented with an auditory stimulus of one frequency ranging between 4-38 kHz at 70 dB sound pressure level (SPL). Mice were trained to lick left in response to tones of one specific ‘center’ frequency (varied between 11-16 kHz across individual animals), and to lick right for tones of any other ‘non-center’ frequency in a 2.5s response period (**Fig. 1a**, **Extended Data** Fig. 1a,b, lick rates were significantly higher in the 1 second following tone offset, N=6 mice). There were eight non-center tones with four tones up to 1.5 octaves lower than the center frequency, and four tones up to 1.5 octaves higher than the center frequency in the full training set (**Fig. 1b**).

**Figure 1.**
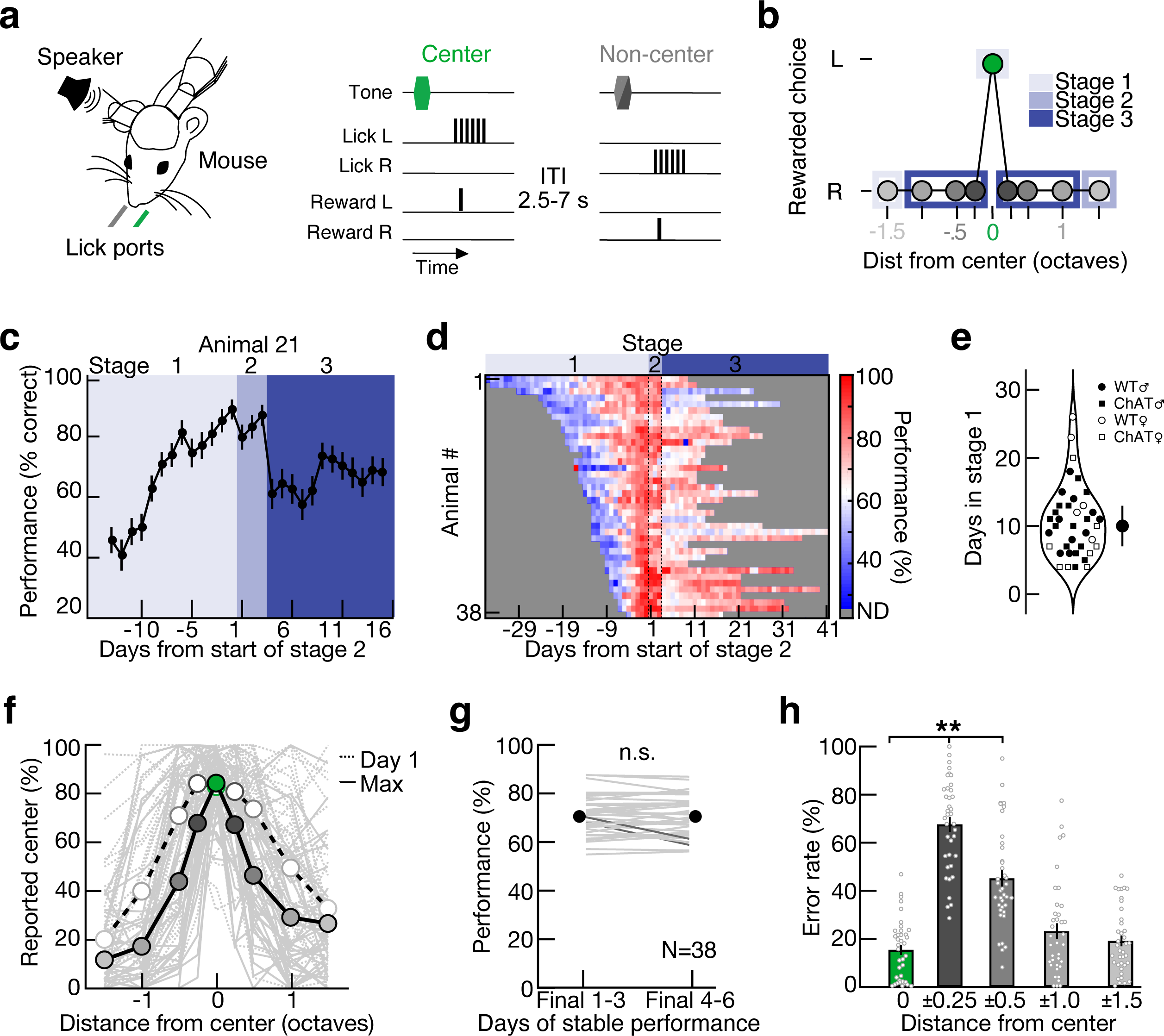
2AFC task for mouse auditory frequency discrimination. **a**, Behavioral schematic showing trial structure. On each trial, a head-restrained and water-restricted mouse is presented with a pure tone of a single frequency for 0.25 seconds at 70 dB SPL. Mice were trained to classify tones as either the center frequency (green) or non-center (shades of gray, with dark gray indicating closest to the center tone frequency and light gray indicating furthest from the center) by licking left (for center) or right (for non-center) in a response epoch 0.25-2.5 seconds from tone presentation. If tones were classified correctly, reward was delivered on the corresponding lick port. Inter-trial interval (ITI), 2.5 to 7 seconds. **b**, Set of auditory training stimuli. The center frequency (11.3-16.0 kHz, specified per animal) was rewarded if the animal licked left; non-center frequency (±0.25-1.5 octaves from center frequency) was rewarded if the animal licked right. A single non-center frequency either -1.5 or 1.5 octaves from center tone was used in stage 1, the other ±1.5 octave frequency was added in stage 2, all other stimuli added in stage 3. **c**, Mean performance across all three stages for example wild-type male mouse. When performance reached ≥80% for three consecutive days in stage 1 (lightest blue), this animal was transitioned to stage 2 (middle blue) for three days before moving to stage 3. Error bars, binomial confidence interval. **d**, Performance for all animals relative to day 1 of stage 2 (N=38 mice). Heat map, % correct performance for each stage of training. Gray, no data (ND) as animals were not trained on those days. **e**, Number of days in stage 1 for all animals (median 10 days, inter-quartile range 7-13 days. Open circles, wild-type females (N=5); filled circles, wild-type males (N=11); open squares, ChAT-Cre females (N=8); filled squares, ChAT-Cre males (N=14). **f**, Percent reported center (i.e., licked left) across all stimuli in on day one of stage 3 (dotted lines) and day of maximum performance (solid lines). Gray lines, individual mice. Colored circles, means (N=38). **g**, Stability of individual performance at end of stage 3 (before starting VNS). Mean overall performance was stable for final six days of stage three; final days 1-3 (performance: 70.5±1.3% correct, mean±s.e.m, N=38) compared to final days 4-6 (performance: 70.5±1.3%, p=0.998 compared to final days 1-3, Student’s two-tailed paired t-test). Individually, only 2/38 animals showed a significant difference (p<0.05, Student’s two-tailed paired t-test) on final days 1-3 vs final days 4-6 (bold lines), 36/38 animals had individually stable performance (thin lines). n.s., not significant. **h**, Error rate for each stimulus. Small circles, individual mice. Error rate was significantly higher at ±0.25 and 0.5 octaves (dark gray, ±0.25 octave error rate: 67.6±3.1%, mean±s.e.m; medium gray, ±0.5 octave error rate: 45.2±3.3%) compared to center (green, center tone error rate; 15.5± 2.1%, p<10^-5^ compared to ±0.25- 0.5, one-way ANOVA with Tukey’s multiple comparisons correction). **, p<0.001.

Mice learned the task in three stages, with additional frequencies added as individual performance improved. During stage 1 (first shaping stage), we only presented two frequencies (the ‘center’ tone and one ‘non-center’ tone 1.5 octaves from center; **Fig. 1b**, light blue). Animals completed stage 1 after reaching our performance criterion (≥80% correct) for three consecutive days (**Fig. 1c**). The number of days to reach performance criteria in stage 1 was variable across individual mice, from 5-37 days (**Fig. 1d,e**). There were no significant differences in performance across stage one by sex or genotype (wild-type vs ChAT-Cre in C57Bl/6J background; wild-type males: 10.8±3.8 days in stage 1, mean±s.d., N=11; wild-type females: 16.4±7.7 days, N=5; ChAT-Cre males: 10.1±3.9 days, N=14; ChAT-Cre females: 7.8±5.4 days, N=8; p=0.05 for genotype, p=0.62 for sex, two-way ANOVA with Tukey’s multiple comparisons correction). Following stage 1, we then introduced one other ‘non-center’ frequency that was 1.5 octaves away from the center frequency in the opposite direction (**Fig. 1b**, medium blue), and trained animals in this second shaping stage for three days (**Fig. 1c,d**). We then introduced all other frequencies (stage 3; **Fig. 1c,d**, N=38 mice), with all animals being trained in stage 3 for at least nine days (**Extended Data** Fig. 1c). Overall performance plateaued at days 7-9 of stage 3, but was variable across individuals (**Extended Data** Fig. 1d,e**,f**, peak performance was not significantly correlated with total days trained in stage 3, Pearson’s R=0.31, minimum five days of training prior to baseline; Days 0-2: 64.7±1.0%; Days 6-8: 70.0±1.2%; Max day ± 1 day: 72.6±1.3%, Mean±s.d.; p<10^-4^ one-way ANOVA with Tukey’s multiple comparisons correction, N=38 animals). Performance improved across non-center frequencies throughout stage 3 with the largest improvement occurring at the frequencies ±0.5 octaves from center (**Fig. 1f**, N=38 mice, day 1 of stage 3 compared to maximum performance, minimum of 3 days of stable performance, **Extended Data** Fig. 1g,h).

Task performance varied considerably across animals even after days to weeks of training (**Fig. 1f, Extended Data** Fig. 1e,f**,g,h**). On the day of peak performance, animals correctly identified frequencies as the center frequency at 84.5±12.7% of the time (N=38 animals, mean±s.d., minimum 3 days of stable performance). Animals generally had stable performance across days after at least nine days in the full version of the task (**Fig. 1g**). Even after performance stabilized, most animals continued to make a significant number of errors on the non-center frequencies closest to the center, either a quarter octave or a half octave away from the center frequency (**Fig. 1h**; error rates at ±0.25 octaves: 67.6±19.0%; error rates at ±0.5 octaves: 45.2±20.6%, mean±s.d.; significantly higher errors compared to center tone responses, p<10^-5^, one-way ANOVA with Tukey’s multiple comparisons correction).

We explored if there was a discrimination threshold further by overtraining a subset of mice for 36.3±12.4 days in stage three (**Extended Data** Fig. 2a, range: 21-48 days, N=6). We then altered the reward structure of the task, such that all errors had a larger impact on the amount of water the mice received. Specifically, we increased the magnitude of reward for all trials (from ∼3µL to ∼6µL) and reducing the overall number of trials (from 400 to 200) for an additional 12-13 days of behavior (**Extended Data** Fig. 2b). There was no significant improvement in performance at the center frequencies or frequencies ±0.25 and ±0.5 octaves from center when comparing all days with the increased reward size and the five finals days with smaller rewards (**Extended Data** Fig. 2c,d**,e,** at 0: small reward: 78.9±4.1% correct, mean±s.e.m., large reward: 83.5±3.3%, p=0.15, Student’s two-tailed paired t-test; at ±0.25: small reward: 36.9±6.8%, large reward: 35.4±4.3%, mean±s.e.m.; p=0.67, Student’s two-tailed paired t-test; at ±0.5: small reward: 64.2±7.0% correct, large reward: 71.4±6.3%, mean±s.e.m.; p=0.08, Student’s two-tailed paired t-test). These data show that while head-fixed mice have relatively stable frequency discrimination abilities, performance remained imperfect even after many days of continued positive reinforcement to correctly resolve frequencies within 0.25-0.5 octaves^35,58,59^.

### Long-term vagus nerve stimulation in mice

Our behavioral results showed that, once a certain level of performance was obtained, additional training did not consistently refine perceptual abilities in an individual animal beyond a certain level. This is consistent with a substantial literature on psychophysics and perceptual learning in humans and other species^5,16–18,35^. We wondered if this was a fixed perceptual limit (e.g., set by physical limitations of the sensory epithelium) or if it could be improved by engagement of central mechanisms of neuromodulation and plasticity. Methods such as transgene expression and optogenetic stimulation can lead to lasting behavioral gains in rodents^21^, but these approaches are currently infeasible or impractical in many species including human subjects. However, stimulation of the vagus nerve has recently been shown to influence neural responses in the central nervous system, altering physiological and cognitive variables^41,43,54,55,60^. Some of these effects may be due to indirect activation of central modulation, but little is known about the underlying mechanisms by which VNS initiates prolonged enhancement of behavior.

We adapted the bipolar VNS cuff used in rats^43^ to make custom vagus nerve cuff electrodes for use in mice^47–49^ and verified successful stimulation of the vagus nerve by measuring impedance of the cuff electrode. We implanted custom cuff electrodes around the left vagus nerve in adult mice (**Fig. 2a**). We measured cuff impedance daily in awake animals for days to months post-implantation over the course of behavioral training (**Extended Data** Fig. 3a,b). Cuff impedances <10 kΩ were considered potentially viable for VNS, and many animals had stable cuff impedances in this range for weeks (**Extended Data** Fig. 3b). We used change in heart rate as a physiological signature for effective VNS (**Extended Data** Fig. 3c-f). When cuff impedance values were very high (e.g., ∼100 MΩ; **Extended Data** Fig. 3d), VNS was ineffective at affecting physiological variables such as heart rate, consistent with our previous work^47^. We thus defined as sham implantations either those electrodes for which impedance values were over 10 kΩ and/or cases in which VNS did not elicit significant changes in heart rate (**Extended Data** Fig. 3f, 7/10 animals with viable cuffs <10kΩ had significantly lower heart rate distributions in sessions with 500 ms of VNS every 10s than in sessions without VNS).

**Figure 2.**
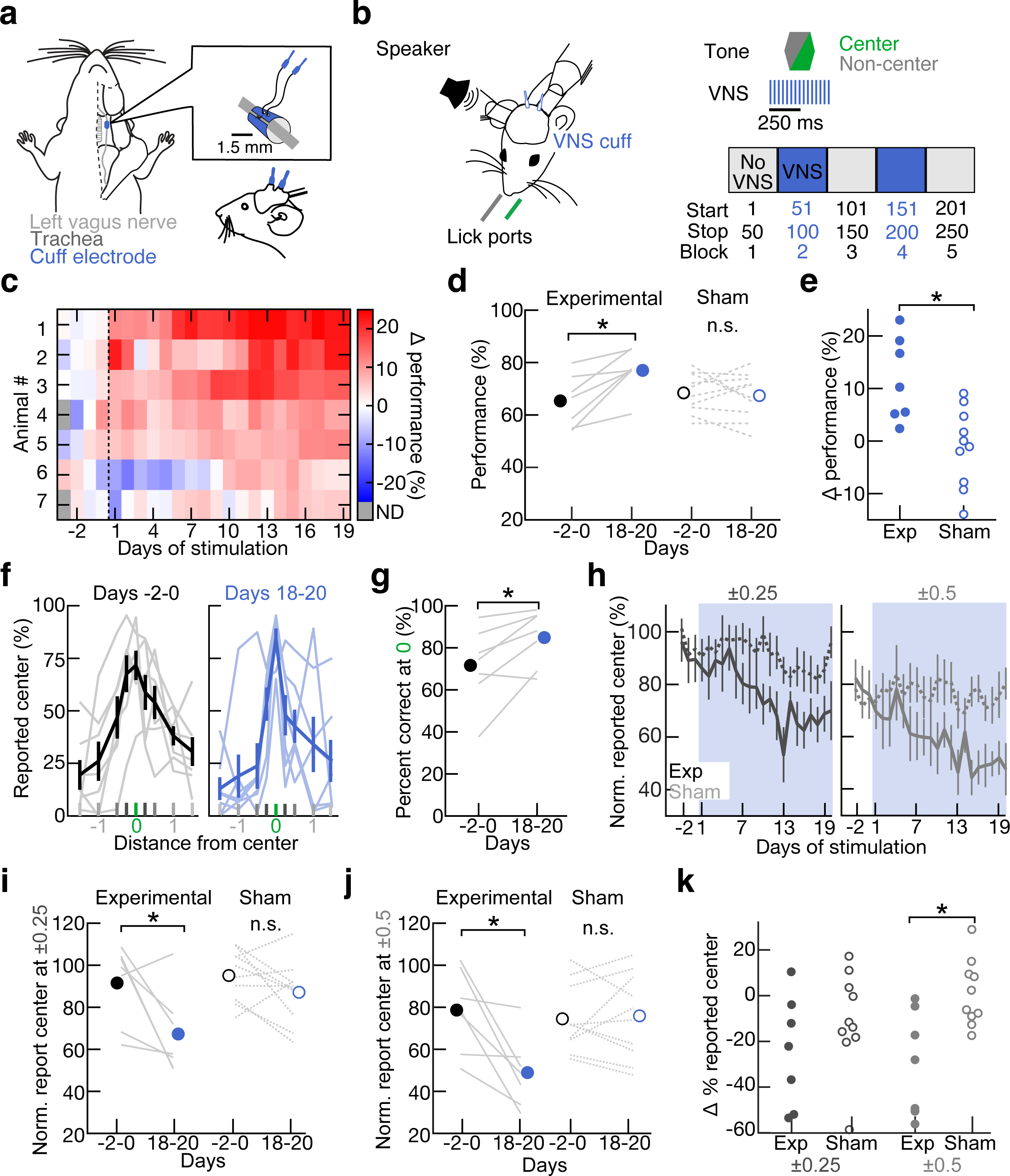
VNS during behavior improves perceptual discrimination over days. **a**, Schematic of cuff electrode implantation on mouse left vagus nerve. **b**, Schematic of VNS pairing during behavior. VNS was performed in two blocks of 50 trials (blue, blocks 2 and 4), interleaved between blocks of no stimulation (gray, blocks 1, 3 and 5). VNS parameters: 500 ms duration, 30 Hz rate, 0.6-0.8 mA intensity, centered around the tone for that trial. During blocks of VNS, stimuli of all frequencies (center and non-center) were presented during behavior. **c**, VNS pairing during behavior gradually improved performance relative to baseline (days after VNS cuff implantation prior to start of VNS pairing) over days (N=7 mice). **d**, Performance over all stimuli improves after VNS pairing, but not in sham control animals. Percent correct on final days of VNS pairing (Days 18-20) across all frequencies in experimental animals (blue, closed circles, 77.1±3.1%, mean±s.e.m., N=7) in comparison to the behavior three days prior to VNS (black, closed circles, ‘Day -2-0’, 65.4±3.8%, p=0.008, Student’s two-tailed paired t-test). Percent correct on final days (Days 18-20) across all frequencies in sham control animals (blue open circles, 67.4±2.8%, mean±s.e.m., N=10) in comparison to behavior three days prior to sham pairing onset (black open circles, 68.5±2.5%, mean±s.e.m., N=10, p=0.65, Student’s two-tailed paired t-test). **e**, Difference in performance on day 18-20 of VNS or sham pairing compared to three days prior to pairing onset in experimental animals (closed circles, 11.7±3.0%, mean±s.e.m., N=7) compared to sham control animals (open circles, -1.1±2.4%, mean±s.e.m., N=10, p=0.004, Student’s one-tailed unpaired t-test). **f**, Percent reported center at each frequency relative to center son the three days prior to VNS onset (black, ‘Days -2-0’, gray lines are individual mice) and behavior on the final days of VNS pairing (blue, ‘Days 18-20’, light blue lines are individual animals, N=7 mice). **g**, Percent correct at center frequency significantly increased on final days of VNS pairing (blue, Days 18-20, 84.9±5.0%, mean±s.e.m.) compared to three days prior to pairing onset (black, Days -2-0, 71.6±7.0%, mean±s.e.m., p=0.02, Student’s two- tailed paired t-test). **h**, Average percent reported center at ±0.25 (dark gray) and ±0.5 (medium gray) octaves normalized by performance at center frequency gradually reduced over days of stimulation (shaded blue) in experimental animals (solid line, N=7), but not sham controls (dotted line, N=10). **i**, Percent reported center normalized by performance at center frequency at ±0.25 octave from center was reduced over days prior to VNS pairing onset (black, Days -2-0) compared to the final days of VNS pairing (blue, Days 18-20) in experimental animals (solid lines, closed circles, Days -2-0: 91.6±7.0%, Days 18-20: 67.2±8.1%, mean±s.e.m., p=0.04, Student’s two-tailed paired t-test), but not sham controls (dotted lines, open circles, Days -2-0: 95.3±3.9%, Days 18-20: 87.2±4.7%, mean±s.e.m., p=0.13, Student’s two-tailed paired t-test). **j**, Percent reported center normalized by performance at center frequency at ±0.5 octave was reduced over days prior to VNS pairing onset (black, Days -2-0) compared to the final days of VNS pairing (blue, Days 18-20) in experimental animals (solid lines, closed circles, Days -2-0: 78.8±7.1%, Days 18-20: 49.2±6.8%, mean±s.e.m., p=0.01, Student’s two-tailed paired t-test), but not sham- implanted control animals (dotted lines, open circles, Days -2-0: 74.8±5.1%, Days 18-20: 75.9±6.4%, mean±s.e.m., p=0.81). **k**, Difference in normalized percent reported center at ±0.25 and ±0.5 octave frequencies from center for sham controls (open circles, difference at ±0.25: -8.1±4.9, difference at ±0.5: 1.1±4.6%, mean±s.e.m.) and experimental animals (closed circles, difference at ±0.25: -24.3.1±9.2%, difference at ±0.5: -29.6±8.6%, mean±s.e.m., for ±0.25: p=0.06, for ±0.5: p=0.002, Student’s one-tailed unpaired t-test).

### Vagus nerve stimulation improves auditory perceptual learning over days

We next asked how VNS might affect the performance of fully trained animals on the 2AFC task. After several days of training on stage 3 (average 14.8+/-8.4 days, range 6-36 days), we implanted 17 animals with a vagus nerve cuff. Of the animals implanted, seven trained mice had <10 kΩ cuff impedances and VNS elicited changes in heart rate (**Extended Data** Fig. 3) and were subsequently included and analyzed as part of the experimental group. All other animals were part of the sham-implanted control group (N=10). Experimental and control animals did not have any significant differences in baseline performance, including peak performance, response rate, and learning rate (**Extended Data** Fig. 4a,b**,c,d,e**). After implantation, animals recovered for about a week with *ad lib* water, and training on the 2AFC task was reinstated after 4-10 days of additional water restriction (total days in stage 3: average 19.0±8.5 days, range 10-40 days). Prior to the start of VNS pairing, implanted animals were trained for 3-7 days post-implantation without VNS, to establish a new baseline level of performance and ensure that this was comparable to their pre- implantation behavior (**Extended Data** Fig. 4f, change in performance (experimental): -1.8±1.6% correct, mean±s.e.m., p=0.38, Student’s two-tailed paired t-test, N=7; change in performance (control): 0.4±1.0% correct, mean±s.e.m., p=0.68, Student’s two-tailed paired t-test, N=10 mice; p=0.28 for experimental and control comparison, Student’s two-tailed unpaired t-test). VNS occurred during behavior in a blockwise manner, with a ‘block’ being a set of 50 trials. In blocks 1, 3, and 5 there was no VNS, and training was identical to that previously described in **Figure 1**. In blocks 2 and 4, VNS was performed concurrently with tone presentation, essentially being paired with all center and non-center stimuli. Specifically, we stimulated the vagus nerve for 500 ms at 30 Hz with a 100 µs pulse width centered around the 250 ms tone, such that VNS began 125 ms before tone onset and continued for 125 ms after tone offset (**Fig. 2b**). By design, this meant that VNS pairing was performed irrespective of trial outcome, in contrast to a recent important study of motor learning^50^. For experimental animals, VNS stimulation intensity was 0.6-0.8 mA, fixed per animal. For sham-implanted, control animals, VNS stimulation intensity varied from 0 to 0.8 mA. Animals underwent VNS pairing for 20 consecutive days.

We found that VNS pairing led to enduring improvements in 2AFC task performance for most animals, which emerged gradually over days and largely enhanced correct responses to the ±0.25 and ±0.5 octave flanking non-center tones (**Fig. 2c**). We initially quantified the change in performance across all stimuli over all five training blocks (‘Δ % correct’) as the overall correct rate on each day relative to the average performance on the final three days of training prior to pairing initiation. Given there was a significant increase in response rate in experimental animals relative to control animals (**Extended Data** Fig. 5a,b**,c,d)**, we included no response trials for comparisons between experimental and control animals. The maximum change in 2AFC task performance occurred several days after initiating VNS pairing during training (**Fig. 2c**, peak performance after 15.4±2.8 days, mean±s.d., N=7 mice). We compared behavioral performance on the final three days of VNS pairing (day 18-20) to the baseline behavioral performance on the three days of training just before the start of VNS pairing in experimental and sham-implanted control animals. Behavioral performance significantly increased in experimental animals (**Fig. 2d**, N=7 mice, before VNS: 65.4±2.8% correct over all center/non-center stimuli, including no response trials (day 18-20 of VNS: 77.1±3.1% correct over all stimuli, p=0.008, Student’s two-tailed paired t-test), but not in sham control animals (**Fig. 2d**, N=10 mice, before VNS: 68.5±2.4% correct, day 18-20 of VNS: 67.4±2.8% correct over all stimuli, p=0.65, Student’s two-tailed paired t-test). Experimental animals had a significantly larger improvement in performance than sham animals (**Fig. 2e**, mean difference in performance (experimental): 11.7±3.0% correct, mean±s.e.m; mean difference (sham): -1.1±2.4% correct, mean±s.e.m., N=10, p=0.002, Student’s one-tailed unpaired t-test).

As VNS occurred with presentation of both center and non-center stimuli, we asked if performance at certain frequencies were specifically improved after days of VNS pairing (**Fig. 2f**, all 17 sham and experimental animals individually displayed in **Extended Data** Fig. 5e). We noticed that, on average, behavioral performance improved at both the center frequency (**Fig. 2f,g**, Day -2-0: 71.6±7.0%, mean±s.e.m.; Day 18-20: 84.9±5.0%, p=0.02, Student’s two-tailed paired t-test) and the non-center tones flanking the center (**Fig. 2f**; ±0.25 and ±0.5 octaves from center). To correct for perceptual improvements at the center frequency across animals, we normalized the behavioral performance at individual frequencies on the final days of VNS pairing (days 18-20) and baseline performance (**Fig. 2h-k**, N=7 experimental mice; N=10 control mice). Most of the improvement in behavioral performance came from gradual reductions in errors at non-center frequencies ±0.25 and 0.5 octave from the center frequency in experimental animals compared to control animals (**Fig. 2h**). The reduction of normalized errors, and resulting improved performance, at frequencies ±0.25-0.5 octaves from center (**Fig. 2h-j**, ‘Experimental’, for ±0.25: p=0.04, Student’s two-tailed paired t-test; for ±0.5: p=0.01, N=7) were not observed for ±0.25-0.5 octave stimuli in control animals receiving sham VNS (**Fig. 2h-j**,’Sham’, for ±0.25: p=0.13, Student’s two-tailed paired t-test; for ±0.5: p=0.81, Student’s two-tailed paired t-test, N=10 mice). Additionally, the reduction in errors between the final days of VNS (days 18-20) and the three days immediately prior were significantly reduced in experimental animals relative to control animals at frequencies ±0.5 octaves from center (**Fig. 2k**, for ±0.5: p=0.002, Student’s one-tailed unpaired t-test).

As the behavioral effects of VNS pairing gradually emerged over several days, we wondered if there were also immediate, acute perceptual effects during the paired training sessions, and/or if there were improvements occurring each day from block to block with successive training within a daily session. We first analyzed behavioral differences across blocks with and without VNS pairing, specifically comparing block one and three (without VNS) vs block two and four (with VNS). We focused on these two blocks of trials to capture potential differences, e.g., in motivational state as animals acquired more water rewards. In sessions with blocks of VNS, performance did not selectively improve during VNS pairing blocks compared to unpaired blocks, across all stimuli in experimental (**Fig. 3a**; block one performance without VNS: 79.5±3.2% correct, mean±s.e.m.; block two performance with VNS: 79.1±1.9%,; block three performance without VNS: 77.6±3.4%; block four performance with VNS: 75.8±2.5%; N=7, p=0.78, one-way ANOVA with Tukey’s multiple comparisons correction), or sham-implanted control animals (**Fig. 3a**; open circles, block one performance without VNS: 73.3±2.7% correct, mean±s.e.m.; block two performance with VNS: 65.5±4.6%; block three performance without VNS: 70.9±3.5%; block four performance with VNS: 64.9±4.8%,; p=0.37, one-way ANOVA with Tukey’s multiple comparisons correction; N=10). Similarly, there was no significant change in performance between blocks with and without VNS for experimental and control animals (**Fig. 3b**; difference in experimental animals: -1.1±1.3% lower in sessions without VNS, mean±s.e.m.; difference in control animals: -6.9±3.8% lower in sessions without VNS, p=0.23, Student’s two-tailed unpaired t-test). There also could be rapid changes in motivation, learning, or other aspects of animal state within a session from block to block independent of VNS pairing. To account for this, we compared performance per block in the three initial baseline sessions prior to initiating VNS pairing. We found that performance was stable across all blocks in these baseline sessions (**Fig. 3c**; block one performance: 68.0±4.3% correct, mean±s.e.m.; block two performance: 63.9±3.6% correct; block three performance: 63.1±3.7% correct; block four performance: 63.7±3.4%; N=7, p=0.78, one-way ANOVA with Tukey’s multiple comparisons correction). Additionally, there was no significant change in performance across specific frequencies in blocks with or without VNS in experimental or control animals (**Fig. 3d,e**, p=0.28 for frequencies, p=0.11 for experimental group, two-way ANOVA with Tukey’s multiple comparisons correction). There was also no significant change in response rate across blocks in both experimental and control animals (**Fig. 3f,g**; experimental means: block one response rate: 97.1±2.0%; block two: 98.2±1.6%; block three: 95.0±3.0%; block four: 95.5±2.8%; N=7, p=0.77, one-way ANOVA with Tukey’s multiple comparisons correction; control means: block one response rate: 96.7±1.6%; block two: 91.9±3.8%; block three: 93.9±2.1%; block four: 85.7±5.7%; N=10, p=0.20, one-way ANOVA with Tukey’s multiple comparisons correction).

**Figure 3.**
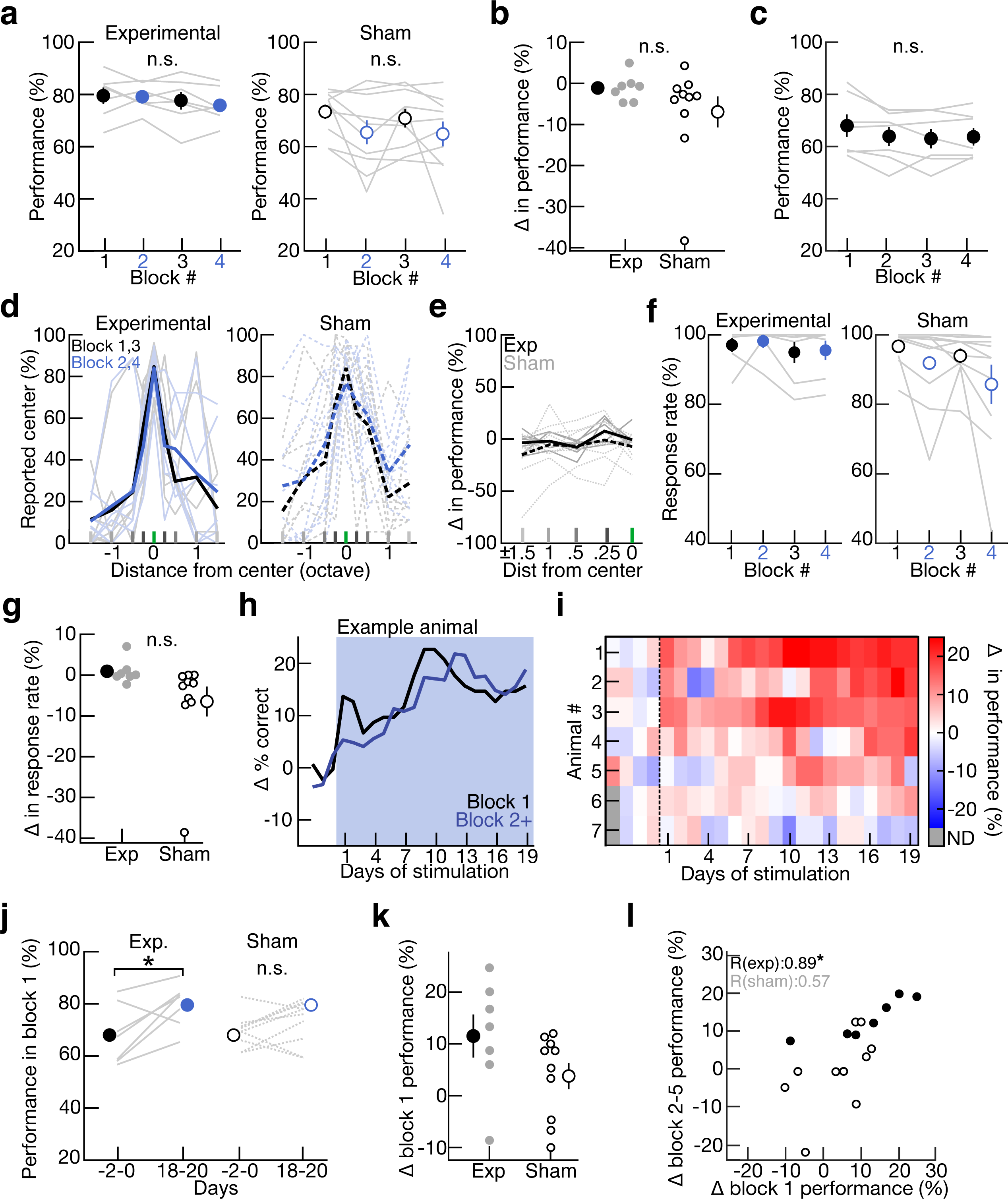
VNS induces long-lasting behavioral improvements. **a**, Performance for all stimuli is stable across blocks 1-4 on final three days with VNS in experimental animals (closed circles, block 1: 79.5±3.2% correct, mean±s.e.m., no VNS, black; block 2: 79.1±1.9%, VNS, blue; block 3: 77.6±3.4%, no VNS, black; block 4: 75.8±2.5%, VNS, blue; p=0.78, one-way ANOVA with Tukey’s multiple comparisons correction; N=7) and control animals (open circles, block 1: 73.3±2.7% correct, mean±s.e.m., no VNS, black; block 2: 65.5±4.6%, VNS, blue; block 3: 70.9±3.5%, no VNS, black; block 4: 64.9±4.8%, VNS, blue; p=0.37, one-way ANOVA with Tukey’s multiple comparisons correction; N=10). **b**, Difference in performance across blocks with and without VNS for experimental (closed circles, average difference: -1.1±1.3%, mean±s.e.m.) and control animals (open circles, average difference: -6.9±3.8%, p=0.23, Student’s two-tailed unpaired t-test). **c**, Performance for all stimuli was stable across blocks 1-4 prior to any days of VNS (block 1: 68.0±4.3% correct, mean±s.e.m., no VNS, black; block 2: 63.9±3.6%, VNS, blue; block 3: 63.1±3.7%, no VNS, black; block 4: 63.7±3.4%, VNS, blue; p=0.78, one-way ANOVA with Tukey’s multiple comparisons correction; N=7). **d**, Percent reported center at each frequency relative to center stimulus on final days of VNS pairing in blocks with VNS (blue, blocks 2 and 4, light blue lines are individual animals) or without (black, blocks 1 and 3, gray lines are individual animals) in experimental (solid lines) and control mice (dotted lines). **e**, Difference in performance across block types (with or without VNS) at each frequency relative to the center stimulus for experimental (solid line) and control animals (dotted line, p=0.28 for frequencies, p=0.11 for experimental group, two-way ANOVA with Tukey’s multiple comparisons correction). **f**, Response rate is stable across block 1-4 on final three days with VNS pairing in experimental (closed circles, block 1: 97.1±2.0% correct, mean±s.e.m., no VNS, black; block 2: 98.2±1.6%, VNS, blue; block 3: 95.0±3.0%, no VNS, black; block 4: 95.5±2.8%, VNS, blue; p=0.77, one-way ANOVA with Tukey’s multiple comparisons correction; N=7) and sham animals (open circles, block 1: 96.7±1.6% correct, mean±s.e.m., no VNS, black; block 2: 91.9±3.8%, “VNS”, blue; block 3: 93.9±2.1%, no VNS, black; block 4: 85.7±5.7%, “VNS”, blue; p=0.20, one-way ANOVA with Tukey’s multiple comparisons correction; N=10). **g**, Difference in response rate across blocks with and without VNS for experimental (closed circles, average difference: 0.9±1.1%, mean±s.e.m.) and control animals (open circles, average difference: -6.5±3.7%, p=0.13, Student’s two-tailed unpaired t-test). **h**, Mean change in percent correct relative to the three baseline days for all stimuli in block one (i.e., 50 trials prior to receiving VNS during behavior that day, black) and all subsequent trials (including 100 trials with VNS, blue) in an example animal. **i**, Summary of change in percent correct for all stimuli in block one (N=7 mice). **j**, Performance in block one during baseline sessions prior to VNS onset (black, ‘Days -2-0’) compared to performance on final days of VNS pairing (blue, ‘Days 18-20’) in experimental (closed circle, solid lines, average performance (Days -2-0): 68.0±4.3%, mean±s.e.m., average performance (Days 18-20): 79.5±3.2%, p=0.03, Student’s two-tailed paired t-test, N=7) and control animals (open circle, dotted lines, average performance (Days -2-0): 69.5±1.9%, mean±s.e.m., average performance (Days 18-20): 73.3±2.7%, p=0.17, Student’s two-tailed paired t-test, N=10). **k**, Difference in performance in block 1 between the three days prior to VNS pairing onset (Days -2-0) and the final three days of VNS pairing (Days 18-20) in experimental (closed circles, average difference: 11.5±4.1%, mean±s.e.m.) and control animals (open circles, average difference: 3.8±2.6%, p=0.057, Student’s one-tailed unpaired t-test). **l**, Correlation between change in performance in block one and all subsequent trials in experimental animals (closed circles, Pearson’s R=0.89, p=0.008, N=7) and control animals (open circles, Pearson’s R=0.57, p=0.09, N=10).

Instead of immediate gains in performance during or just after VNS pairing, these results indicate that behavioral changes took days to emerge, accruing over daily sessions with VNS pairing. This would suggest that the gradual changes in performance might be observed even in block one across days, before any VNS pairing occurred on that day. To test this hypothesis, we compared performance on block one (i.e., the first 50 2AFC trials without VNS pairing of each day) to performance over all blocks across days. Experimental animals exhibited similar rates of improvement over time for block one alone compared to performance on all trials following block 1 (**Fig. 3h-j**). Across animals, the change in performance during block one of the final days of VNS was significantly correlated with change in performance over all other trials (**Fig. 3l**, Pearson’s R=0.78, p=0.008). In contrast, control animals did not exhibit the same significant improvements in performance in block one (**Fig. 3j-l**).

### VNS pairing enables long-term cortical plasticity

Our results demonstrate that VNS pairing enhanced perceptual learning even beyond the limits achieved just by behavioral training. As these gains emerged gradually over days, we suspected that a mechanism related to enduring neuroplasticity was being engaged by VNS pairing. Previous studies in rats pairing pure tones with VNS^43,51^ found changes to neuronal populations and tonotopic maps of primary auditory cortex (A1), strongly suggesting that auditory cortex is a locus of potentially behaviorally-relevant plasticity.

To identify possible long-term changes in cortical responses and receptive fields triggered by VNS pairing, we performed longitudinal two-photon imaging in mouse auditory cortex of seven untrained animals (**Fig. 4a**; 6 wild-type females, 1 wild-type male). We injected all mice with a CaMKII-GCaMP6f in auditory cortex, waited 2-4 weeks for viral expression, and then implanted a VNS cuff electrode on the left vagus nerve. We paired a tone of a single frequency with VNS by coincidental presentation every 2.5 seconds for 5 minutes each day for at least 5 days (up to 20 days). We performed these experiments outside the context of behavior, so that we could monitor neural changes that might be more directly attributed to VNS-mediated plasticity, avoiding the confound of cortical plasticity that might occur due to training^8,9,11,12^. In each animal, we tracked a consistent population of neurons over days of pairing (**Fig. 4a-c**, **Extended Data** Fig. 6). We also presented pure tones of 4-64 kHz every 1-3 days throughout the pairing procedure to monitor changes across a wider frequency range of the receptive fields (**Fig. 4a**). Additionally, after the initial pairing session, we presented the same set of frequencies both 15 minutes and 2 hours after VNS pairing to look for more acute changes of auditory responses in a subset of animals. In control animals (1 wild-type female, 3 wild-type males), we performed the same pairing protocol, presenting a tone of a single frequency every 2.5 seconds, but without coincident VNS (**Fig. 4a**). Changes in tuning were assessed in the same manner.

**Figure 4.**
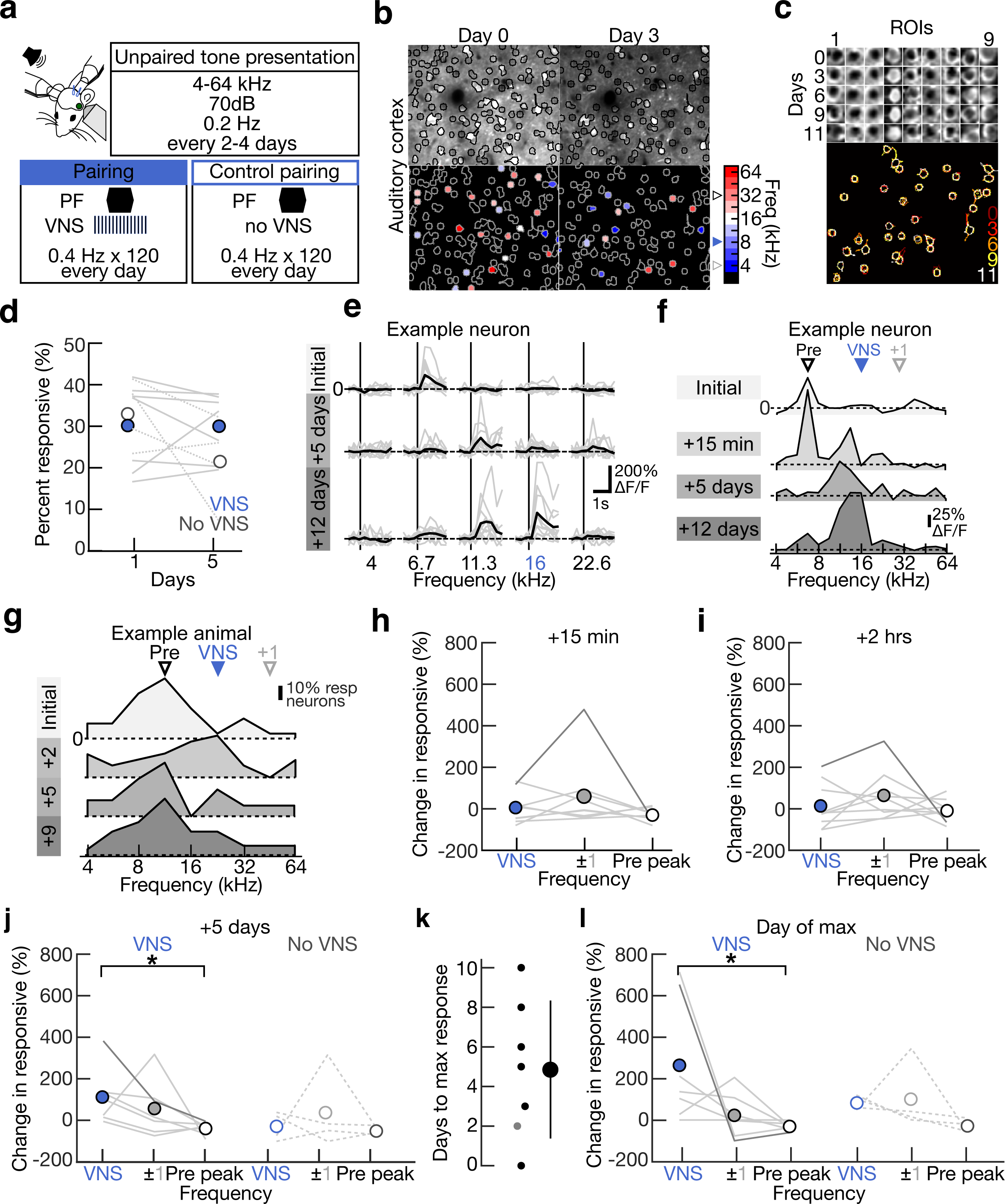
VNS enables long-term plasticity in excitatory neurons in auditory cortex. **a**, Schematic of VNS-tone pairing while performing two photon imaging of excitatory neurons in auditory cortex. During VNS-tone pairing, VNS was for 500ms at 30Hz, centered around the 250ms tone of one frequency every 2.5s. Pairing was done every day for up to 20 days. Control animals also heard a tone of one frequency at the same rate without the coincident VNS. During unpaired, passive tone presentation, tones ranging from 4-64 kHz at 70 dB are presented for 250ms every 5-10s. Passive tone presentation was performed every 1-3 days. **b**, Portion of imaging region with ROIs colored for the frequency eliciting the maximal response in example animal during two days of passive tone presentation (Day 0: prior to receiving any VNS-tone pairing; Day 3: after receiving three days of VNS-tone pairing). Initial best frequency (“Pre-peak”, black, open arrow), pairing frequency (“P”, blue, closed arrow) and frequency ± one octave from the paired frequency (“±1”, gray, open arrow) represented for example animal on color bar. **c**, Example ROIs tracked across 5 days of passive tone presentation (top: subset of neurons tracked over time; bottom: spatial overlap of all neurons tracked across all sessions within consistent imaging area, different colors represent different days of imaging). **d**, Percentage of significantly responsive neurons over two time points (initial day of unpaired passive tone presentation and day 5 of VNS pairing) in control (dotted lines, open circles; initial day: 33.0±8.1% significantly responsive to any frequency, mean±s.d, day 5: 21.5±10.6% responsive, N=4 mice, n=192±35.8 neurons per animal) and experimental animals (solid lines, closed circles; initial day: 30.2±10.7% significantly responsive to any frequency, mean±s.d, day 5: 30.0±7.9% responsive, N=7 mice, n=145.3±20.5 neurons per animals, mean±s.d., p=0.96 for experimental animals, p=0.20 for sham animals, Student’s paired t-test, p=0.11 for difference across time points for experimental and sham animals, Student’s unpaired t-test). **e**, Example neuron during initial passive tone presentation, after 5 days of VNS-tone pairing and after 12 days of VNS-tone pairing for select frequencies. The paired tone was 16 kHz (seen in blue) for this animal. Mean response for each frequency is represented in black and individual trials are shown in gray. **f**, Passive tuning curves across all frequencies for example neuron prior to first VNS pairing (lightest gray), +15 minutes post first VNS pairing (light gray), 5 days of VNS pairing (medium gray), and after 12 days of pairing (dark gray). ‘VNS’, paired tone was 16 kHz for this animal. ‘+1’, tone one octave from paired tone. ‘Pre’, initial best frequency pre-peak across the population. **g**, Percentage of neurons responsive to 4 to 64 kHz prior to VNS pairing (lightest gray), 3 days of VNS pairing (light gray), 5 days of VNS pairing (medium gray) and after 9 days of pairing (dark gray) in example animal. The paired tone was 22.7 kHz for this animal. **h**, Change in percent responsive 15 minutes after initial VNS-tone pairing relative to initial responses at the frequency with the maximum initial response (Pre-peak; open circle; -31.4±29.9%; mean±s.d.), paired frequency (blue; 4.4±75.7%; mean±s.d.) and frequency one octave away from the paired frequency (gray; 59.3±164.1%; mean±s.d., N=9 mice, n=1303 neurons, 144.8±33.2 neurons per animal, mean±s.d.; p=0.21; one-way ANOVA with Tukey’s multiple comparisons correction). Example animal from **g** highlighted in dark gray. **i**, Change in percent responsive 2 hours after initial VNS-tone pairing relative to initial responses at the frequency with the maximum initial response (Pre peak; open circle; - 9.0±52.9%; mean±s.d.), paired frequency (blue; 14.1±110.0%; mean±s.d.) and frequency one octave away from the paired frequency (gray; 64.8±119.5%; mean±s.d., N=9 mice, n=1303 neurons, 144.8±33.2 neurons per animal, mean±s.d.; p=0.29; one-way ANOVA with Tukey’s multiple comparisons correction). Example animal from **g** highlighted in dark gray. **j**, Change in percent responsive at day 5 of pairing relative to initial responses for at the frequency with the maximum initial response (‘Pre peak’, black), paired frequency (blue), and frequency one octave away from the paired frequency (±1, gray) for experimental (paired: 113.3±132.2%, mean±s.d.; Pre peak: -38.4±29.4%; Freq ±1: 57.9±133.4%; N=7 mice, n=953 total neurons, 136.1±16.5 neurons per animal; p=0.055; one-way ANOVA with Tukey’s multiple comparisons correction) and sham animals (paired: -28.0±68.5%, mean±s.d.; Pre peak: -50.4±19.0%; Freq ±1: 38.2±188.9%; N=4 mice, n=737 neurons total, 184.3±37.1 neurons per animal; p=0.56; one-way ANOVA with Tukey’s multiple comparisons correction). Example animal from **g** highlighted in dark gray. **k**, Distribution of days to maximum response at paired frequency (4.9±3.5 days; mean±s.d.; N=7 mice). Gray circle is animal represented in **g. l**, Change in percent responsive at day of maximum population response to paired frequency relative to initial response at with the maximum initial response (‘Pre peak’, black), paired frequency (blue), and frequency one octave away from the paired frequency (±1, gray) for experimental (paired: 263.2±295.2%, mean±s.d.; Pre peak: -31.3±19.4%; Freq ±1: 22.3±105.1%; N=7 mice, n=934 neurons total, 133.4±15.5 neurons per animal; p=0.02; one-way ANOVA with Tukey’s multiple comparisons correction) and sham animals (paired: 82.0±33.6%, mean±s.d.; Pre peak: - 28.7±30.1%; Freq ±1: 100.1±163.8%; mean±s.d., N=4 mice, n=750 neurons total, 187.5±43.8 neurons per animal; p=0.19; one-way ANOVA with Tukey’s multiple comparisons correction). Day of maximum population response was determined for individual animals in **k**. Example animal from **g** highlighted in dark gray.

We imaged from layer 2/3 excitatory neurons in each animal (N=7 mice, 144.8±33.2 neurons per animal, mean±s.d.) in a 300 μm x 300 μm region. 455/1081 neurons (42.1%) recorded prior to VNS-tone pairing were responsive to at least one frequency between 4-64 kHz, and 371/953 neurons (37.8%) recorded on day 5 of pairing were responsive (with 465 cells imaged on both sessions). There was no change in the percentage of significantly responsive neurons across days and type of pairing (sham or experimental pairing) (**Fig. 4d,** p=0.96 for experimental animals, p=0.20 for sham animals, Student’s paired t-test, p=0.11 for difference across time points for experimental and sham animals, Student’s unpaired t-test). For each animal, we computed the overall best frequency across auditory cortex in the region of imaging (initial best frequency, ‘Pre peak’), and 78/455 responsive neurons (17.1%) on the first imaging day had significant or maximal responses to that local best frequency. We then chose a frequency that was initially under- represented at the level of individual neuronal tuning profiles as the ‘paired frequency’, with 29/455 responsive neurons (6.4%) initially responding to that frequency across animals before VNS pairing.

VNS pairing modified the tuning curves of auditory-responsive cells over minutes to days, as quantified by the percentage of active neurons responding to a certain stimulus (**Fig. 4e-g**). We measured the change in number of responsive neurons at three frequencies: the initial best frequency (‘Pre peak’; black), the paired frequency (‘P’, blue), and a frequency one octave from the paired frequency (‘±1’, gray) to ask how stimulus-specific these changes were. We focused our analysis of frequency receptive fields to several time points throughout pairing, trying to highlight any plasticity that could occur over minutes to hours (**Fig. 4h,i**) or over days (**Fig. 4j-l**). Since there was significant individual variability in reaching peak behavioral improvement, we postulated that a similar variability could be present in VNS-mediated long-term plasticity across animals. To that end, we looked at the number of days it took animals to show maximal changes at the paired frequency (**Fig. 4k,l**).

At shorter time scales (15 minutes post initial pairing, **Fig. 4h**, 2 hours post initial pairing, **Fig. 4i**), there were not significant changes in response to the paired frequency (‘P’, blue), initial best frequency (‘Pre peak’, black), or the frequency one octave away from the paired frequency (±1, gray). However, consistent with the gradual time course of behavioral enhancement we observed with VNS, reliable and lasting cortical changes required several days to emerge. At both long-term time points (after five days of pairing and the day with maximal changes at the paired frequency), two major changes resulted from VNS pairing but not control pairing (**Fig. 4j-l**). First, the overall number of neurons responding to the paired frequency significantly increased (Day 5: 113.3±132.2% increase at paired frequency, mean±s.d., one-way ANOVA with Tukey’s multiple comparisons correction; response at paired frequency was significantly different than ‘Pre peak’ with p<0.05; Max: 263.2±295.2% increase; mean±s.d., response at paired frequency was significantly different than ‘Pre peak’ with p=0.02, one-way ANOVA with Tukey’s multiple comparisons correction, N=7 mice). Second, the number of neurons responding to the original best frequency decreased (Day 5: -38.4±29.4% decrease, mean±s.d.; Max: -31.3±19.4% decrease). The maximum change in percent of responsive neurons relative to initial tuning at the paired frequency occurred after several days of pairing (4.9±3.5 days; mean±s.d.; n=1018 neurons initially; n=934 neurons on day of max response; N=7 mice). As in our past studies of the behavioral effects of pairing auditory stimuli with basal forebrain stimulation^35^, we hypothesize that both the changes to the paired and original best frequencies are required together, in order to effectively reshape cortical tuning curves for improving sensory perception.

### VNS activates cholinergic basal forebrain neurons projecting to auditory cortex

A major outstanding question regarding the utility of VNS across species is the mechanisms of action. Specifically, it remains unclear how and where VNS leads to either direct or indirect activation of different brain regions (including auditory cortex). We reasoned that central neuromodulation was involved, given the diverse set of clinical applications of VNS in human subjects^39–42^, the slower rates for consistent perceptual improvement and cortical plasticity seen in mice ^35,61^ and the known anatomical projections from NTS^56,57^. However, many of the effects of VNS have been historically attributed to the titratable activation of the locus coeruleus and consequent central noradrenergic release^49,54,55,62^. The potential contribution of cholinergic modulation to VNS has been more controversial. Conventionally the cholinergic basal forebrain was thought to receive only indirect input from NTS via locus coeruleus^56,57^. While behavioral evidence links motor learning with the recruitment of cholinergic basal forebrain neurons via VNS^50,63^, there are differences in the functional organization of cholinergic inputs to various cortical regions^30,64–67^. Thus, we next wanted to establish if the central cholinergic system might contribute to the sensory perceptual improvements we observed with VNS.

We asked if basal forebrain cholinergic cells were functionally activated by VNS. To do this, we performed fiber photometry of basal forebrain cholinergic neurons in untrained animals with VNS cuffs. We injected ChAT-Cre mice with a Cre-dependent GCaMP6s in basal forebrain and implanted a fiber above the injection site. VNS was applied every 10 seconds for 20 trials and lasted 500 ms at 30Hz at 0.8-1.0 mA (**Fig. 5a**). Consistent with other results^50^, we found 500 ms of VNS was sufficient to induce prolonged activation of basal forebrain cholinergic neurons in a subset of trials (**Fig. 5b,c**). The average activation of cholinergic cell bodies by VNS on individual trials was negatively correlated with baseline fluorescence, such that trials with relatively high baseline prior to VNS onset were more likely to have lower average activation (**Extended Data** Fig. 7a,b, Pearson’s R=-0.30, p=0.02).

**Figure 5.**
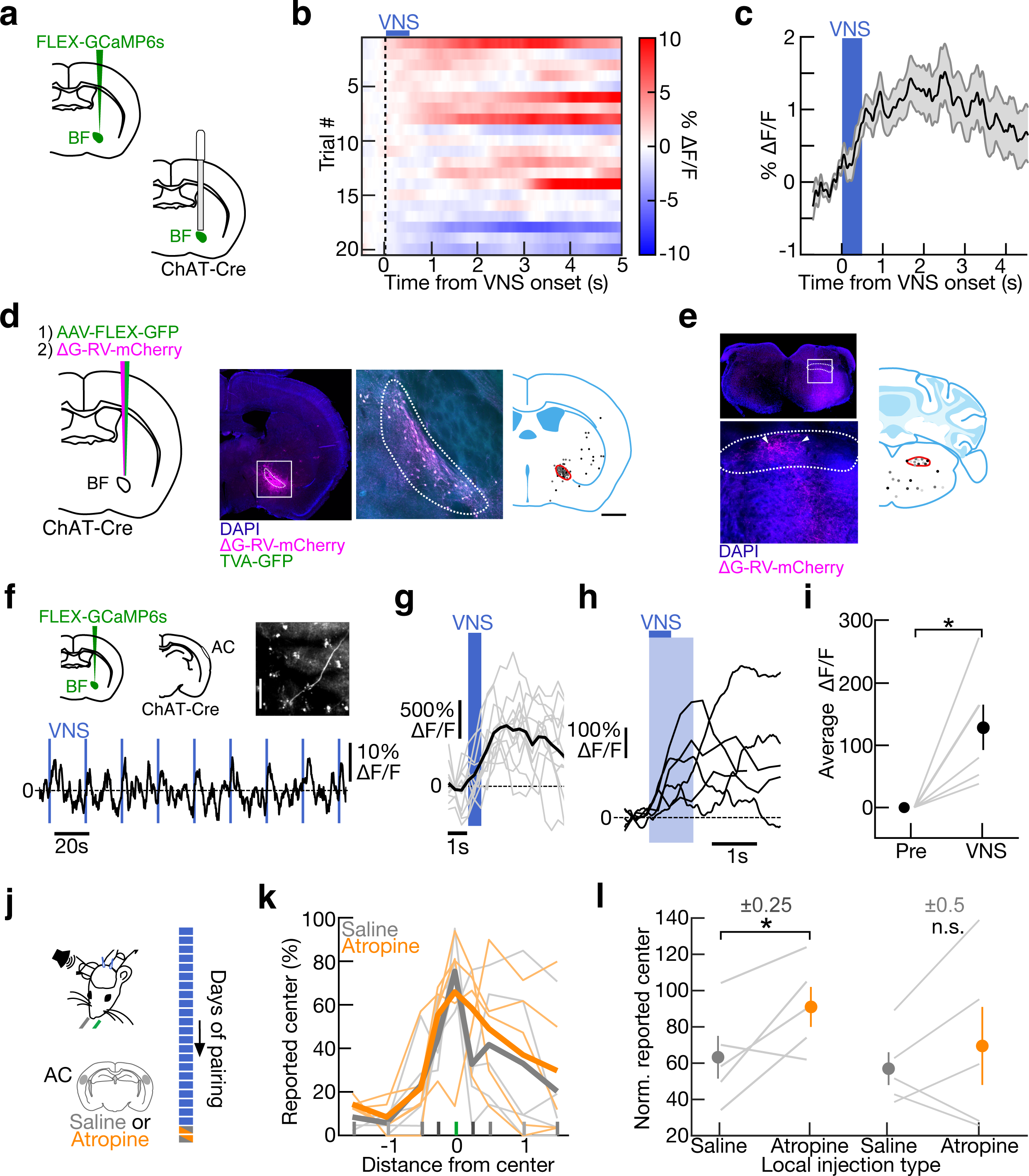
Stimulating the vagus nerve activates auditory-cortical projecting cholinergic basal forebrain. **a**, Schematic of fiber photometry from cholinergic basal forebrain neurons. **b**, ΔF/F from all VNS applications for one example animal. VNS was applied for 500 ms at 30 Hz at 1.0 mA every 10 seconds. **c**, Average ΔF/F for all VNS trials across animals with mean baseline ΔF/F less than zero (N=3 mice; n=36 trials). **d**, Schematic for mapping inputs to cholinergic basal forebrain neurons using retrograde, Cre- dependent, pseudotyped monosynaptic rabies. Example image of injection area in basal forebrain, corresponding section from the Allen Brain Atlas, and location of GFP labeled neurons. The regions of interest are highlighted in red. Each animal is represented in a distinct shade of gray (N=3). **e**, Location of mCherry labeled neurons in section containing NTS labeled by dots in shades of gray. Regions of interest are highlighted in red. Each animal is represented in a distinct shade of gray (N=6). **f**, Schematic of imaging of basal forebrain cholinergic axons in auditory cortex. Trace of fluorescent activity is from one example animal from one example region. VNS applied for 500 ms at 30 Hz at 0.8 mA every 20 s. **g**, VNS triggered average from one example region. Example animal is the same shown in **f**. **h**, Distribution of time to max ΔF/F relative to VNS onset across all VNS applications in all regions in all animals (N=6 mice, 16.5±6.3 regions per animal, mean±s.d., n=270 trials). VNS was applied for 500ms at 30Hz at 0.8-1.0 mA every 2.5-26.7s. **i**, Average ΔF/F for 1s baseline prior to VNS onset and 1s following VNS onset (shaded in light blue in **l**, N=6, VNS: 128.7±36.4% ΔF/F, mean±s.e.m., p=0.02, Student’s two-tailed paired t-test). **j**, Schematic for local injection of either atropine (orange) or vehicle (gray, control) in auditory cortex after 20 days of VNS pairing during behavior. After local injection, VNS was applied during behavior in the same manner as described in Figure 2. **k**, Percent reported center across all frequencies for either atropine (individual animals represented in light orange, mean in darker orange) or vehicle (individual animals represented in light gray, mean in darker gray). **l**, Normalized percent reported center for atropine or vehicle at frequencies ±0.25 (atropine: orange, 91.0±24.5%, mean±s.d.; vehicle: gray, 63.3±26.4%, mean±s.d., N=5, p=0.03, Student’s one-tailed paired t-test) and ±0.5 from center (atropine: orange, 69.5±48.0%, mean±s.d.; vehicle: gray, 57.0±20.2%, mean±s.d., N=5, p=0.21, Student’s one-tailed paired t-test).

Since VNS robustly activated cholinergic basal forebrain neurons, we wanted to better understand the anatomical pathway contributing to this activation. Previous work has shown that NTS projects to LC^56^, which in turn projects to basal forebrain^57^. We wanted verify if cholinergic neurons of the basal forebrain were primarily indirectly activated by VNS via NTS projections to the locus coeruleus or whether there was another previously underexplored pathway. To do this, we performed retrograde tracing studies from cholinergic neurons in basal forebrain using a Cre-inducible, retrograde pseudotyped monosynaptic rabies virus^68,69^. We targeted cholinergic neurons by injecting pAAV-TREtight-mTagBFP2-B19G and pAAV-syn-FLEX-splitTVA-EGFP-tTA in the basal forebrain of ChAT-Cre mice (N=3) or a AAV1 EF1a-FLEX-GTB (N=3) followed by an injection of SADΔG-mCherry two weeks later (**Fig. 5d**). We found mCherry expression in both NTS and locus coeruleus, indicating that basal forebrain cholinergic neurons likely receive direct input from both areas (**Fig. 5e**, **Extended Data** Fig. 7c,d). We verified this result by injecting a Cre-dependent tdTomato with a retrograde promoter in basal forebrain of TH-Cre animals (**Extended Data** Fig 7e). We again saw tdTomato expression in NTS (**Extended Data** Fig. 7e). These anatomical studies show that cholinergic basal forebrain neurons receive input from TH+ neurons in both locus coeruleus and NTS (**Extended Data** Fig. 7f).

We next wanted to explore if VNS could activate basal forebrain cholinergic axons projecting to auditory cortex. Previous studies found that VNS can activate cholinergic fibers in cortex^48,49^, but for much longer stimulation periods than used in our task (1-10 second periods of VNS). Thus, we used two-photon imaging of cholinergic axon fibers in auditory cortex (**Fig. 5f**) to more specifically monitor recruitment of cortically-projecting cholinergic inputs that might produce the behavioral and neuronal effects of VNS tone pairing described above (VNS for 500 ms at 30 Hz at 0.8 mA). ChAT-Cre mice were injected with Cre-dependent GCaMP6s in basal forebrain, a window was implanted over auditory cortex and a VNS cuff was applied to the left vagus nerve (N=6). VNS was applied for 500 ms at 30 Hz at 0.8-1.0 mA every 2.5-26.6 seconds. We found that VNS strongly activated many cholinergic axon fibers in auditory cortex on a trial-by-trial basis at various timescales with some trials reaching maximum activation during or immediate after VNS offset and some persisting for several seconds (**Fig. 5g**). VNS activated cholinergic axons across regions within auditory cortex and across animals (**Fig. 5h,i**). Interestingly, unlike with activation of cholinergic cell bodies via VNS, there was no significant correlation between baseline activity and VNS evoked activation (**Extended Data** Fig. 7g, Pearson’s R=0.04, p=0.71). The difference in trial-by-trial activation of axons and cell bodies could be due to differences in experimental techniques (measurement of bulk activation vs activation of axon segments) and/or due to the diversity of inputs and projection targets of subsets of cholinergic neurons^64,65^.

We then asked if VNS-mediated task improvement required acetylcholine receptors in auditory cortex acutely. To do this, after we infused either atropine (a muscarinic acetylcholine receptor antagonist) or vehicle in auditory cortex 30 minutes prior to behavior after animals underwent 20 days of VNS pairing (**Fig. 5j**). We then applied VNS in the same manner as described previously (block-wise for 50 trials on/off, 500ms centered around center and non-center frequencies). Blocking activation of local acetylcholine receptors increased the error rate at the frequencies ±0.25 octaves from center (**Fig. 5k,l**, at ±0.25: atropine: orange, 91.0±24.5%, mean±s.d.; vehicle: gray, 63.3±26.4%, mean±s.d., N=5, p=0.03, Student’s one-tailed paired t-test). Therefore, VNS applied at levels that induce behavioral improvements activates auditory cortical projecting cholinergic neurons in basal forebrain and acetylcholine in auditory cortex impacts behavior acutely.

### Optogenetic manipulation of the central cholinergic system and VNS

In the final set of experiments, given the activation of cholinergic fibers in auditory cortex, we asked how cholinergic modulation related to long-term VNS pairing in behaving animals. Since areas of cortex receive input from distinct subsets of cholinergic neurons^64,65^, we aimed to target only the auditory cortical-projecting cholinergic neurons. For projection-specific opsin expression, we bilaterally injected a retrograde, Cre-dependent channelrhodopsin-2 (ChR2) virus in auditory cortex and implanted optic fibers above basal forebrain in eight ChAT-Cre animals trained on the 2AFC task (**Fig. 6a, Extended Data** Fig. 8a). After 9+ days in stage three, instead of VNS pairing (as these animals were not cuffed), we optogenetically stimulated auditory cortical-projecting cholinergic neurons in the same manner as VNS: 500 ms duration, 30 Hz pulse rate, centered around the 250 ms tone during training blocks two and four. Control animals were trained in the same manner, did not have significantly different baseline behavior (**Extended Data** Fig. 8b,c,) and underwent the same pairing protocol as optogenetic stimulation animals but were injected with a retrograde, Cre-dependent fluorophore virus (rg-FLEX-tdTomato), instead of channelrhodopsin-2. Given that there was not a significant change in response rate before and after optogenetic pairing (**Extended Data** Fig. 8d), no response trials were not included in comparisons for control and ChR2-positive animals.

**Figure 6.**
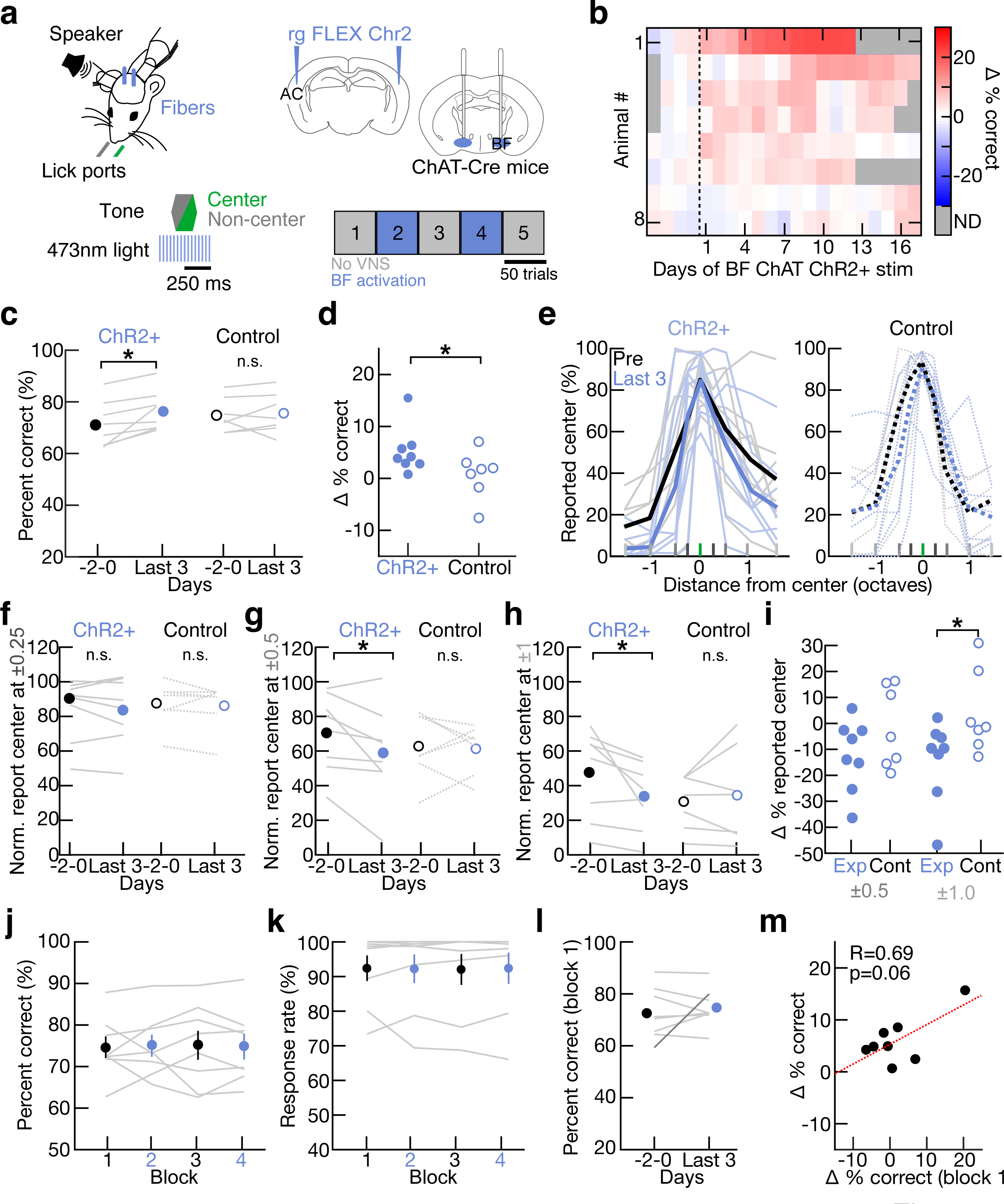
Activating auditory cortical projecting cholinergic basal forebrain neurons improves perceptual performance. **a**, Schematic of optogenetic activation of auditory cortical projecting cholinergic BF neurons during behavior. Auditory cortical projecting cholinergic BF neurons were targeted using a retrograde Cre-dependent channelrhodopsin injected into auditory cortex of ChAT-Cre mice. ChAT+ BF activation was applied in the same blockwise fashion as previously described for 500 ms at 30 Hz centered around the tones. **b**, Optogenetic activation of ChAT+ BF neurons during behavior gradually improved performance over days (N=8 mice). **c**, Performance over all stimuli improves after optogenetic pairing in experimental animals (closed circles), but not control animals (open circles). Percent correct on the final three days of channelrhodopsin pairing for each animal all frequencies (light blue; experimental mean: 76.3±2.8%, mean±s.e.m., N=8; control mean: 75.6±3.0%, N=7) in comparison to the behavior three days prior to pairing onset (black; experimental mean: 71.1±3.1%, p=0.01, Student’s two-tailed paired t-test, N=8; control mean: 74.8±2.5%, p=0.67, Student’s two-tailed paired t-test, N=7). **d**, Difference in performance on the last three days of channelrhodopsin pairing relative to three days prior to pairing onset for experimental (closed circles, 5.3±1.6%, mean±s.e.m., N=8) or control animals (open circles, 0.8±1.7%, N=7, p=0.04, Student’s one-tailed unpaired t-test). **e**, Percent reported center at each frequency relative to center stimulus on the three days prior to optogenetic pairing onset (black) and behavior on the last three days of channelrhodopsin pairing (light blue) in experimental (solid lines, N=8) and control animals (dotted lines, N=7). **f**, Percent reported center normalized by performance at the center frequency at ±0.25 octave frequency from center was not significantly reduced over days prior to channelrhodopsin pairing onset (black, Days -2-0) compared to the final days of channelrhodopsin pairing (blue, ‘Last 3’) in experimental animals (solid lines, closed circles, Days -2-0: 84.6±6.4%, Last 3: 83.6±6.9%, mean±s.e.m., p=0.67, Student’s two-tailed paired t-test) or control animals (dotted lines, open circles, Days -2-0: 87.5±4.8%, Last 3: 86.2±5.0%, mean±s.e.m., p=0.56, Student’s two-tailed paired t-test). **g**, Percent reported center normalized by performance at the center frequency at ±0.5 octave frequency from center was significantly reduced over days prior to optogenetic pairing onset (black, Days -2-0) compared to the final days of channelrhodopsin pairing (blue, ‘Last 3’) in experimental animals (solid lines, closed circles, Days -2-0: 70.9±7.9%, Last 3: 58.9±9.7%, mean±s.e.m., p=0.04, Student’s two-tailed paired t-test), but not control animals (dotted lines, open circles, Days -2-0: 62.9±7.2%, Last 3: 61.3±5.3%, mean±s.e.m., p=0.80, Student’s two-tailed paired t-test). **h**, Percent reported center normalized by performance at the center frequency at ±1 octave frequency from center was not significantly reduced over days prior to channelrhodopsin pairing onset (black, Days -2-0) compared to the final days of channelrhodopsin pairing (blue, ‘Last 3’) in experimental animals (solid lines, closed circles, Days -2-0: 47.5±9.0%, Last 3: 33.5±6.8%, mean±s.e.m., p=0.04, Student’s two-tailed paired t-test) or control animals (dotted lines, open circles, Days -2-0: 30.7±5.9%, Last 3: 34.4±10.2%, mean±s.e.m., p=0.55, Student’s two-tailed paired t-test). **i**, Difference in normalized percent reported center at ±0.5 and ±1 octave frequencies from center for control animals (open circles, difference at ±0.5: -1.6±5.8, difference at ±1: 3.8±5.9%, mean±s.e.m.) and experimental animals (closed circles, difference at ±0.5: -12.0±4.8%, difference at ±1: -14.0±5.5%, mean±s.e.m., for ±0.5: p=0.09, for ±1: p=0.02, Student’s one-tailed unpaired t-test). **j**, Performance for all stimuli is stable across blocks 1-4 on final three days with optogenetic stimulation in experimental animals (closed circles, block 1: 74.7±2.6% correct, mean±s.e.m., no ChR2+ pairing, black; block 2: 75.1±2.7%, ChR2+ pairing, blue; block 3: 75.2±3.5%, no ChR2+ pairing, black; block 4: 74.8±3.1%, ChR2+ pairing, blue; p=0.99, one-way ANOVA with Tukey’s multiple comparisons correction; N=8). **k**, Response rate is stable across block 1-4 on final three days with optogenetic pairing in experimental animals (block 1: 92.3±3.7% correct, mean±s.e.m., no ChR2+ pairing, black; block 2: 92.2±4.1%, ChR2+ pairing, blue; block 3: 92.0±4.5%, no ChR2+ pairing, black; block 4: 92.3±4.5%, ChR2+ pairing, blue; p=0.99, one-way ANOVA with Tukey’s multiple comparisons correction; N=8). **l**, Performance in block one during baseline sessions prior to optogenetic pairing onset (black, Days -2-0: 72.6±3.5%, mean±s.e.m.) compared to performance on final three days of channelrhodopsin pairing (blue, Last 3: 74.7±2.6%, p=0.50, Student’s two-tailed paired t-test, N=8). **m**, Correlation between change in performance in block one and all subsequent trials (Pearson’s R=0.67, p=0.06, N=8).

We found that optogenetic activation of cholinergic neurons during behavior led to similar improvements in perceptual learning as with VNS pairing. The maximum change in 2AFC task performance occurred several days after optogenetic pairing (**Fig. 6b**, N=8 mice). We compared behavioral performance on the last three days optogenetic pairing (‘Last 3’) to three days of baseline behavioral performance (‘-2-0’). Maximum performance was significantly increased past the level obtained just from days of stage three training in optogenetic stimulation animals (**Fig. 6c,d**, before optogenetic pairing: 71.1±3.1% correct over all center/non-center stimuli, mean±s.e.m, last three days of pairing: 76.3±2.8% correct over all stimuli, N=8 mice, p=0.01, Student’s two-tailed paired t-test), but not control animals (**Fig. 6c,d**, pre: 74.8±2.5% correct, mean±s.e.m, last three days of pairing: 75.6±3.0%, N=7, p=0.67, Student’s two-tailed paired t-test; difference in performance prior to optogenetic activation and during, experimental: 5.3±1.6%, mean±s.e.m., N=8, control: 0.8±1.7%, N=7, p=0.04, Student’s one-tailed unpaired t-test). Performance gains occurred at specific non-center flanking frequencies as with VNS pairing, specifically at ±0.5-1.5 octaves from center (**Fig. 6e**, behavior of individual animals shown in **Extended Data** Fig. 8e). Optogenetic stimulation animals, but not control animals, had a significant reduction in errors at the frequencies ±0.5 and ±1 octave from the center frequency, but not ±0.25 octaves from center (**Fig. 6f,g,h**,**i**, control: difference at ±0.25: -1.4±2.2, difference at ±0.5: -1.6±5.8%, difference at ±1: 3.8±5.9, mean±s.e.m.; optogenetic stimulation animals: difference at ±0.25: -1.0±2.2%, difference at ±0.5: -12.0±4.8%, difference at ±1: -14.0±5.5%, mean±s.e.m., for ±0.25: p=0.55, for ±0.5: p=0.09, for ±1: p=0.02, Student’s one-tailed unpaired t-test). Similar to VNS experimental animals, optogenetic activation of cholinergic basal forebrain neurons did not acutely improve performance (**Fig. 6j**) or alter response rate (**Fig. 6k**). While behavioral performance in block 1 did not show the significant improvement over time seen in VNS experimental animals (**Fig. 6l**), the change in block 1 performance and overall performance on the last three days of pairing relative to baseline were positively correlated (**Fig. 6m**, R=0.67, p=0.06, N=8), consistent with VNS experimental animals.

While the decrease in error rates with optogenetic pairing occurred over a broader range of stimulus frequencies than VNS pairing, the time course and specific reductions to off-center flanking tones was a consistent feature of both pairing methods. These observations support the hypothesis that the predominant effects of VNS pairing for this 2AFC task in mice is through cholinergic recruitment. To test this hypothesis more directly, in **Figure 7** we show the results of optogenetic suppression of cholinergic basal forebrain neurons during VNS pairing. In other trained ChAT-Cre animals (N=6 mice also shown in **Fig. 1**), we bilaterally injected a Cre-dependent inhibitory opsin (archaerhodopsin) in the basal forebrain, implanted optic fibers bilaterally over basal forebrain (**Extended Data** Fig. 9a), and implanted a cuff electrode on the left vagus nerve (**Fig. 7a**). We verified successful cuff implantation in six animals by looking for both low impedance and reliable reduction in heart rate in response to VNS (**Extended Data** Fig. 9b,c**,d**). After 9+ days of stage three training, we began VNS pairing as before, i.e., performing VNS on each training day during blocks two and four. During VNS pairing blocks, we concurrently optogenetically suppressed ChAT+ basal forebrain neurons continuously for 500 msec starting 125 ms before tone onset. Given that there was not a significant change in response rate before and after optogenetic pairing (**Extended Data** Fig. 9e), no response trials were not included in comparisons for control and ChR2-positive animals. Optogenetic suppression of basal forebrain cholinergic neurons completely prevented behavioral gains from VNS pairing in overall performance (**Fig. 7b,c**, N=6 mice, before VNS and optogenetic suppression: 62.2±2.3% correct over all center/non-center stimuli, day 18-20 of VNS and optogenetic suppression: 62.7±2.3% correct over all stimuli, p=0.79, Student’s two-tailed paired t-test) and compared to performance gains seen in experimental animals (**Fig. 7d**, p=0.01, Student’s two-tailed unpaired t-test). Inhibition of cholinergic basal forebrain neurons during VNS also prevented VNS-mediated improvements at frequencies ±0.25 and ±0.5 octaves from center (**Fig. 7e-g**, individual animals shown in **Extended Data** Fig. 9f). No long-term changes were induced in these animals, as performance on block one remained similar to baseline even after 18-20 days of VNS when cholinergic activity was suppressed (**Fig. 7h,i**). Collectively our results highlight the importance of the central cholinergic modulatory system for enhancements of perceptual learning via VNS.

**Figure 7.**
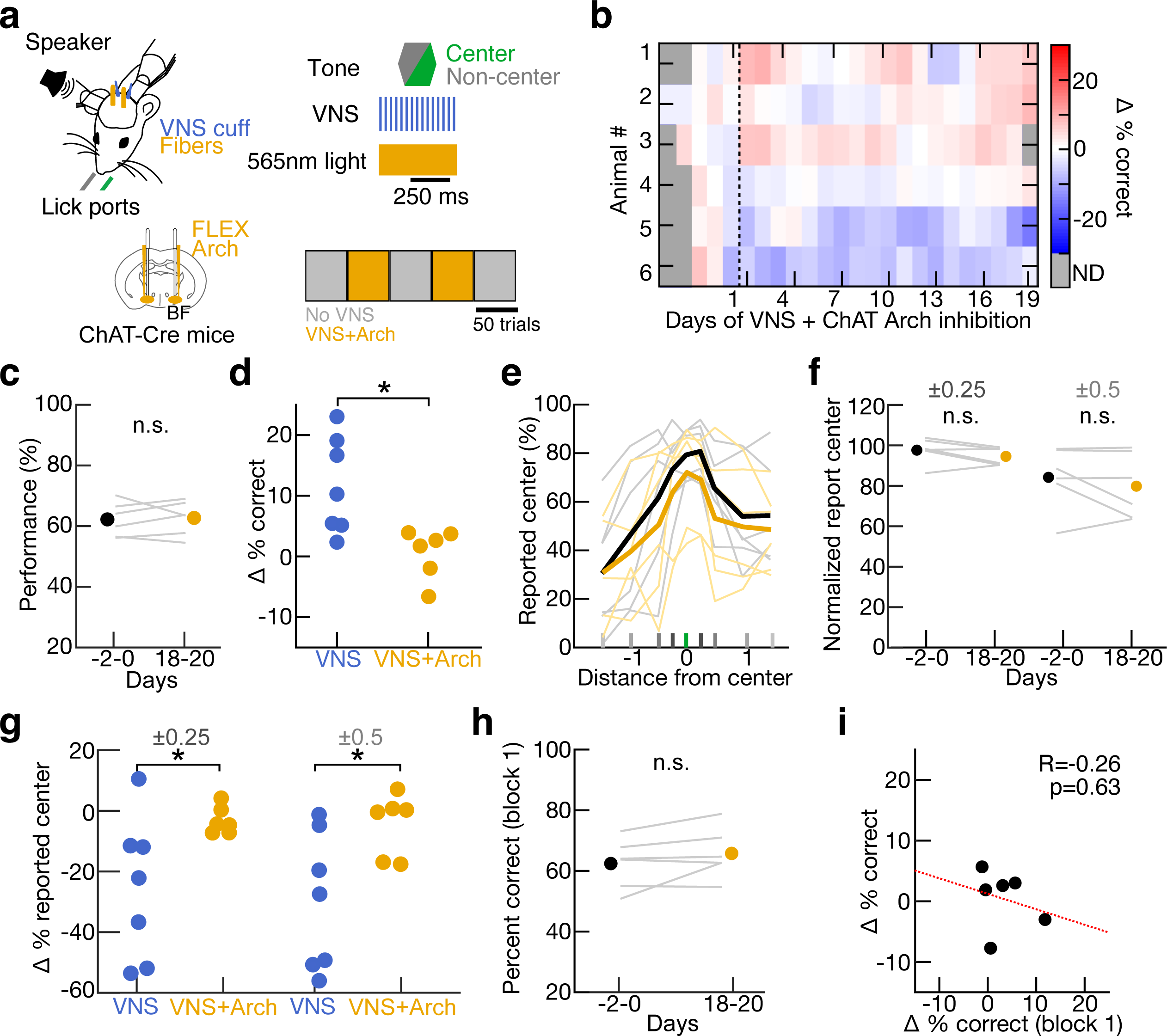
Inhibiting cholinergic basal forebrain neurons during VNS blunts VNS- mediated perceptual improvements. **a**, Schematic of VNS and optogenetic inhibition of cholinergic BF neurons during behavior. VNS was applied in the same blockwise fashion as previously described. Cholinergic BF neurons were optogenetically inhibited using a Cre-dependent Archaerhodopsin and green light (565 nm) for the entire duration of VNS during behavior (500 ms). **b**, Optogenetic inhibition of ChAT+ BF neurons during VNS during behavior abolishes improved performance (N=6 mice). **c**, Performance did not improve after optogenetic inhibition during VNS. Percent correct on the final three days of optogenetic inhibition during VNS (Days 18-20; yellow; 62.7±2.3%, mean±s.e.m., N=6) in comparison to the behavior three days prior to VNS (black; 62.2±2.3%, p=0.79, Student’s two-tailed paired t-test). **d**, Difference in performance on day 18-20 of VNS pairing relative to three days prior to pairing onset for animals receiving VNS with optogenetic inhibition of cholinergic basal forebrain neurons (yellow, 0.5±1.7%, mean±s.e.m., N=6) or without (blue, 11.7±3.0%, mean±s.e.m., N=7, p=0.01, Student’s two-tailed unpaired t-test). **e**, Percent reported center at each frequency relative to center stimulus on the three days prior to start of optogenetic inhibition with VNS pairing onset (black) and the final three days of pairing (yellow, Days 18-20, N=6 mice). **f**, Percent reported center normalized by performance at the center frequency at ±0.25 and ±0.5 octave frequency from center was not significantly reduced over days prior to VNS pairing onset (black, Days -2-0 (±0.25): 97.6±2.5%, Days -2-0 (±0.5): 84.3±6.3%, mean±s.e.m.) compared to the final days of VNS pairing (yellow, Days 18-20 (±0.25): 94.5±1.8%, Days 18-20 (±0.5): 79.8±6.5%, p(±0.25)=0.15, p(±0.5)=0.33, Student’s two-tailed paired t-test). **g**, Difference in normalized percent reported center at ±0.25 and ±0.5 octave frequencies from center for animals with optogenetic inhibition of cholinergic basal forebrain neurons during VNS pairing (yellow, difference at ±0.25: -3.1±1.8%, difference at ±0.5: -4.5 ±4.2%, mean±s.e.m.) or without (blue, difference at ±0.25: -24.3±9.2%, difference at ±0.5: -29.6 ±8.6%, mean±s.e.m., for ±0.25: p=0.04, for ±0.5: p=0.03, Student’s one-tailed unpaired t- test). **h**, Performance in block one during baseline sessions prior to pairing onset (black, Days -2-0: 62.4±3.4%, mean±s.e.m.) compared to performance on final days of VNS and optogenetic inhibition of cholinergic basal forebrain neuron pairing (yellow, Days 18-20: 65.8±3.4%, p=0.15, Student’s two-tailed paired t-test, N=6). **i**, Correlation between change in performance in block one and all subsequent trials (Pearson’s R=-0.26, p=0.63, N=6).

## Discussion

Although the central nervous system is highly plastic, there are limitations on the extent of changes induced by different experiences or mechanisms. For the auditory 2AFC task we used here, animals made the most errors within half an octave of the ‘lick left’ reference tone; VNS pairing could reduce error rates and sharpen behavioral performance, albeit not completely. Some limits in terms of plasticity and behavioral improvement are set by the physical properties of the sensory epithelium. In the visual domain, this includes retinal photoreceptor density and single-photon sensitivity^70,71^; for the auditory system, the ability to resolve different frequencies is constrained by the biophysics of the cochlear membrane and hair cell dynamics^58,59^. These properties are likely to provide hard bounds on perceptual resolution and are difficult to overcome by alternative mechanisms for potential plasticity.

Other limitations, however, might be due to other factors such as motivational state, behavioral engagement, and/or the understanding of task rules and variables. These other factors likely reflect the activation (or lack thereof) of central modulatory systems including the cholinergic basal forebrain^31,72,73^. Indeed, here we found that bidirectional regulation of cholinergic modulation could affect 2AFC task performance, with activation of cholinergic neurons enhancing task performance, whereas suppressing cholinergic neuron activity blocked the effects of VNS. Similarly, we and others have previously shown that engagement in similar auditory^29–31,67^ or visual^27^ tasks activates neurons of the cholinergic basal forebrain, that cholinergic modulation is important for task performance^28,31^, and that artificially enhancing cholinergic modulation (via pharmacology or electrical/optogenetic stimulation) can boost task performance past the levels achieved purely by behavioral training^24,28,31,34,35^. It remains unclear why the cholinergic basal forebrain is not normally fully engaged during task performance. As the effects of VNS on task performance were weakest for the best-performing animals, it is possible that the degree of cholinergic activation is a major predictor of individual sensory processing abilities and perceptual learning rates.

The vagus nerve is remarkably complex, connecting to with several peripheral organs to provide a multiplexed input to the brainstem NTS^74^. In turn, the NTS sends projections to several regions important for central neuromodulation, including the oxytocin system of the hypothalamus^75^, the noradrenergic locus coeruleus^56,57,75^, and the cholinergic basal forebrain^64–66^. Potentially any or all of these systems might be activated by VNS, although little is known how patterns of vagal activation (either naturally occurring or via VNS) lead to similar or differential recruitment of these diverse modulatory systems. The specific parameters used for VNS (e.g., stimulation rate or intensity) might also lead to variability in terms of which types of vagus nerve fibers^76^ and/or downstream systems are reliably activated. We used here a consensus VNS parameter set found to generally be effective across a number of different outcome measures and species^43,47–50^, but there could be other stimulation regimes or longer-term dynamics that might shift the net effects of VNS. There may also be species-specificity in terms of central consequences of VNS. While aspects of the neuroanatomical organization of the vagus and NTS seem largely conserved, the thickness and composition of the vagus nerve bundle is considerably different across species, which would result in different subsets of fibers being activated by electrical stimulation^77,78^.

Despite this complexity, it appeared that VNS as used here in mice largely resulted in cholinergic modulation of auditory cortex. Although NTS projects to locus coeruleus^56^ and locus coeruleus projects to basal forebrain^57^, our anatomical tracing studies also revealed direct projections from both the locus coeruleus and NTS to the basal forebrain, indicating that the cholinergic system might be a major point of convergence for vagal inputs from the brainstem to affect cortical function and behavior. Previous studies have shown that enhancement of perceptual learning and behavioral performance with cholinergic modulation can be surprisingly slow, taking days to weeks to emerge^24,61^. This is similar to the gradual improvements in 2AFC performance we observed here with VNS pairing. In contrast, the effects of locus coeruleus pairing can be much more rapid, but possibly at the expense of initial performance^61^, and over-activation of the locus coeruleus can cause behavioral arrest^79^. Previous work investigating how autonomous system can activate central brain areas showed that activation of the autonomous system can cause anxiety-like behaviors potentially via activation of the noradrenergic system^80^. This might account for why we observed essentially no effects of VNS stimulation during or immediately after paired blocks, but instead that the results of VNS pairing required days to be observed across animals. The relatively slow expression of changes after VNS pairing might reflect a sleep-dependent process requiring days for consolidation, and/or plasticity within neuromodulatory areas such as the basal forebrain^29,30^ or locus coeruleus^61^ than then impact enhanced functionality only after reaching some threshold level of modification. Additionally, if these systems are somewhat in opposition or require temporally precise coordination^63,81^, especially in terms of their short-term effects, it is possible that combined stimulation of multiple modulatory centers leads to no net improvement in the moment. Our work provides evidence that the sustained activation of the autonomous system over many days can reverse the initial impact of activation from producing anxiety-like behaviors^80^ to mediating lasting perceptual improvements.

Neuroprosthetic devices can provide successful treatments for a wide range of debilitating conditions, and perhaps could be adopted for use in augmenting performance outside of clinical care. Some types of devices are implanted centrally, such as deep brain stimulation electrodes, temporal lobe electrodes for regulating seizures, or motor cortex implants in cases of tetraplegia. The invasive nature of central implants limits their utility to only the most severe conditions. In contrast, more human subjects have received neuroprosthetic implants for peripheral nerve stimulation; e.g., cochlear implants have been used in over 500,000 people and are largely successful in terms of hearing restoration. Vagus nerve cuffs have been used in over 100,000 human subjects, mainly for treatment of epilepsy as well as other conditions^40^. A new generation of less-invasive or non-invasive peripheral nerve stimulators targeting the hypoglossal nerve or the auricular branch of the vagus nerve may be promising in terms of efficacy, although much more work is required to validate these devices and determine optimal stimulation regimes^82^. Our data suggest that some outcomes of successful VNS might take days, weeks, or even longer to be revealed, and our results provide a potential mechanistic basis by which VNS can enhance auditory perceptual learning.

## Methods

### Animals

All procedures were approved under an NYU Langone Institutional Animal Care and Use Committee protocol. Male and female mice aged 6-20 weeks old were used in all experiments (**Fig. 1**: N=38, 25 male, 13 female; **Fig. 2**: N=17, 12 male, 5 female; **Fig. 3**: N=17, 12 male, 5 female; **Fig. 4**: N=13, 4 male, 9 female; **Fig. 5**. N=14, 7 male, 7 female; **Fig. 6**: N=15, 7 male, 8 female; Fig. 7: N=6, 6 male). Genotypes used were wild-type C57BL/6J (The Jackson Laboratory, Stock No: 000664), ChAT-Cre (The Jackson Laboratory, Stock No: 028861), and TH-Cre (The Jackson Laboratory, Stock No: 008601). All mice had a C57BL/6J background. Mice were housed in a temperature and humidity controlled room maintained on a 12 hour light/dark cycle. Animals used in behavior were given 1 mL water/day. If their weight dropped below 80% of original, they were given *ad libitum* water until weight returned to ≥80% original value.

### 2AFC Behavioral Training

Behavioral events (lick detection, auditory stimulus delivery, water reward delivery) were monitored and controlled by custom MATLAB programs interfacing with an RZ6 auditory digital signal processor (Tucker-Davis Technologies) via RPvdsEx software (Tucker-Davis Technologies). Licks were detected using capacitance sensors (SparkFun, Part number: AT42QT1011) and water was delivered using solenoids (The Lee Company, Part number: LHDA0581215H). Animals were restrained using custom headposts (Ponoko).

Behavioral training on the auditory 2AFC task began after 7+ days of water restriction. Training started with habituation to head-fixation with water delivered to the mouse while it sat in a plexiglass tube. This was followed by lick port sampling sessions, in which the animal could receive water by alternating licking between the two ports with a minimum of 3 seconds between possible rewards. Mice typically learned to alternate ports while licking for 2-4 µL water droplets in 2-4 sessions. Once animals reliably licked to receive water from lick ports, stage 1 training was begun (i.e., animals were trained to lick left for the center frequency and lick right for one non-center frequency). The center frequency was chosen to be either 11.3, 13.4, or 16 kHz (each animal had a single consistent center frequency pseudo-randomly selected from those three values). Non-center frequencies were set per animal to be ±0.25, ±0.5, ±1.0, and ±1.5 octaves from the selected center frequency. In stage 1, the only non-center frequency was either +1.5 octaves or -1.5 octaves from center (and whether higher or lower frequency was also pseudo-randomly assigned per animal).

In stage 1, while an animal’s performance remained <80% correct, they were rewarded with water regardless of behavior choice on 15% of trials to help promote consistent licking during training. Once performance reached ≥80% correct for three consecutive days in stage 1, animals moved to stage 2 in which the other non-center frequency (either ± 1.5 octaves away) was added. After three days in stage 2, animals moved to stage 3 regardless of performance (in which all other non-center stimuli ±0.25, ±0.5, and ±1.0, octaves from the center frequency were also presented and rewarded for right-side licking).

On each trial, a 250 ms tone was presented and animals had to classify the tone as the center frequency (green) or any other frequency (shades of gray). Stimuli were presented at 70 dB SPL in a pseudorandom order, such that the likelihood of center:non-center was 1:1 (with frequency uniformly chosen from the non-center distribution on non-center trials). After a 250 ms delay, animals had to lick left to report the stimulus as ‘center’ and had to lick right to report the stimulus as ‘non-center’. If the animal did not respond during the 2.5 seconds of the response epoch, the trial was classified as a ‘no response’ trial (which were excluded from analysis except where otherwise noted). If the lick response was correct, a small water reward (2-4 µL) was delivered to the corresponding lick port. Inter-trial intervals were 3±0.5 seconds (mean±s.d.) on trials with a correct response and 6±0.5 seconds (mean±s.d.) on trials with an incorrect response or without a response. Animals were not punished for licking outside of the response epoch. Animals generally performed between 350-500 trials/day.

### Vagus nerve stimulation in mice

The custom cuff electrode design was adapted for mice from a previous cuff electrode designed for rats^43,44^. A bipolar stimulating peripheral nerve cuff electrode was custom built using micro-renathane tubing (Braintree Scientific, Part number: MRE-040), coated platinum iridium wire (Medwire, Part number: 10IR9/49T), and gold pins (Mouser Electronics, Part number: 575- 100140). The tubing was used as the base for the portion of the cuff that interacts with the nerve. It was cut in 1.0-1.5 mm segments, with an interior diameter of 0.025 inches to allow for nerve swelling after implantation. Two platinum iridium wires with the coating removed were glued (Dentsply Sirona, Triad gel) to the interior of the tubing about 0.5 mm apart. Gold pins were soldered to the end of the wires opposite the tubing. The coating on the wire was only removed for small portions at both ends to limit non-coated contact only to the gold pins and within the interior circumference of the tubing. Wires were 2.5-3.0 cm in length in order to span from the skull to the position of the cervical vagus. Non-absorbable silk suture string (Braintree Scientific, Part number: SUT-S 104) was added to each side of the cuff opening to improve cuff handling and manipulation during implantation. Viable cuff electrodes were determined by an impedance reading between 1- 10 kΩ at 1 kHz when submerged in saline (Peak Instruments LCR45).

Impedance measurements were taken each day of stimulation (Peak Instruments LCR45, Frequency: 1kHz). Measurements of breathing rates, SpO_2_, and heart rates were collected using a thigh sensor (MouseOx) in animals lightly anesthetized with 0.75-1.5% isoflurane during VNS. The vagus nerve was stimulated using a high-current stimulus isolator (World Precision Instruments A385) triggered by a digital signal processor (RZ6, Tucker-Davis Technologies). VNS parameters were based on previous work^43,48–50:^ 100 µs pulse width, 30 Hz stimulation rate, 0.5 second duration. Stimulation intensity was 0.6-0.8 mA, based on the magnitude of the effect on vitals (VNS intensity was 0.8 mA for imaging studies of **Figs. 4,5**).

Surgeries were performed on mice aged 6-12 weeks. Mice were anesthetized using isoflurane (1.0-2.5%) and positioned in a stereotaxic frame (Kopf, model 923-B). Body temperature was maintained at 37°C with a heating pad and rectal temperature probe. For behavior and imaging experiments, a custom headpost^31^ was attached to the skull using dental cement (C&B-metabond), after thorough cleaning using hydrogen peroxide. Following all surgical procedures, animals were given a nutritionally-fortified water gel for recovery assistance (Clear H_2_O DietGel Recovery, Part number: 72-06-5022).

For cuff implantation, a magnetic retractor base plate (Fine Science Tools, Part number: 1800-03) was used instead of a stereotaxic frame to allow for more flexible positioning of the animal. Mice were positioned semi-supine at a 45° angel on their right side. Hair on the chest was removed with Nair from the sternum to the left shoulder. The surgical site was sterilized by alternating 70% ethanol and betadine washes. A 1.5 cm incision was made 0.5-1.0 cm to the left of the top of the sternum, then the submandibular gland was separated and retracted from connective tissue. Using blunt forceps (Fine Science Tools, Part number: 11231-30), the left sternocleidomastoid and omohyoid muscles were separated and retracted until the carotid sheath was accessible. A sterilized cuff electrode was led subcutaneously from the left side of the scalp incision, between the ear and eye, and down to the chest incision site. Once positioned, the cuff end of the electrode was deposited near the carotid sheath and the gold pin leads remained exposed on the head. Using sharp forceps (Fine Science Tools, Part number: 11251-30), a 4 mm stretch of the cervical vagus nerve was isolated from surrounding nerves and vasculature without direct contact with the nerve. The cuff electrode was positioned around the vagus nerve so the nerve was not taut and the non-coated intra-cuff wires had even contact with the nerve. The cuff was knotted closed with non-absorbable silk suture string and muscles were returned to their original positions. Absorbable sutures (Ethicon, Part number: W1621T) were occasionally made on the muscles to keep the cuff from lifting the nerve ventrally. The submandibular gland was repositioned and the skin was sutured closed with absorbable sutures (Ethicon, Part number: VCP433). The sutures were sterilized with betadine and then sealed with surgical glue (Meridian). The electrode cuff leads were secured near the headpost using dental cement (C&B-metabond). The incision was covered with 4% topical lidocaine (L.M.X. 4). Cuff electrode impedance (1 kHz) was recorded immediately after surgery and maintained a similar measurement from before implantation. Successful cuff implantation was verified using both impedance measurements and changes in vitals readings as described above.

All VNS during behavior started after the animals reached stable performance after a minimum of nine days from start of stage three. All animals in either the VNS or sham group received 17-20 days of stimulation. On each day of stimulation, animals performed a total of 400 trials, with 100 trials (two blocks of 50 trials, blocks two and four) of behavior with VNS. All auditory stimuli were stimulated to avoid the animal using solely VNS to identify specific stimuli.

VNS lasted 500 ms and was centered around the tone, i.e., it started 125ms prior to tone onset and ended 125ms after tone offset. Stimulation days ended with 100 trials of unstimulated behavior.

### Two-photon calcium imaging

Cranial window implantation over left auditory cortex was performed, as previously described^31^. For cell body imaging, 1.0 µL of diluted CaMKII.GCaMP6f (AAV1, diluted 1:3 with dPBS or AAV9, diluted 1:10 with dPBS, Addgene number: 100834) was injected in three locations throughout auditory cortex (1.5 mm from lambda, along lateral suture). For axon imaging, 1.0 µL of pAAV.Syn.Flex.GCaMP6s.WPRE.SV40 (diluted 1:2 in dPBS; Addgene: 100845-AAV5; AP: -0.5 mm, ML: -1.8 mm, DV: -4.5 mm from brain surface) was injected in basal forebrain of heterozygous ChAT-Cre mice.

Two-photon fluorescence of GCaMP6f/s was excited at 900 nm using a mode locked Ti:Sapphire laser (MaiTai, Spectra-Physics, Mountain View, CA) and detected in the green channel. Imaging was performed on a multiphoton imaging system (Moveable Objective Microscope, Sutter Instruments) equipped with a water immersion objective (20X, NA=0.95, Olympus) and the emission path was shielded from external light contamination. Images were collected using ScanImage (Vidrio). To image auditory cortex, the objective was tilted to an angle of 50–60°. Awake animals were head-fixed under the microscope and the stimulus isolator was connected to the VNS cuff. We imaged using one of two systems: either a microscope covering approximately ∼300 μm^2^ regions with galvo-galvo scanning at ∼4Hz (0.26 s/frame) or a microscope covering approximately ∼450 μm^2^ regions with resonance scanning at ∼30Hz (laser power ≤40 mW).

For VNS pairing while imaging excitatory neurons, the speaker was ∼10 cm away from the ear contralateral to the window. A consistent region of excitatory neurons in layer 2/3 of A1 (based on vasculature and relative orientation of neurons) was imaged over all days of pairing. For baseline imaging, pure tones (70 dB SPL, 4–64 kHz, 250 ms, 10 ms cosine on/off ramps, quarter-octave spacing, 10 trials for each frequency) were delivered in pseudo-random sequence every 5 seconds. During pairing, one frequency (chosen based on the initial tuning of the area) was played concurrently with VNS every 2.5 seconds for 5 minutes.

Collected data were processed using the Suite2p analysis pipeline^83^. Recorded frames were aligned using a non-rigid motion correction algorithm. Regions of interest (representing excitatory neurons or cholinergic axon segments) were segmented in a semi-automated manner using a Suite2p based classifier. Additional ROIs were manually drawn on an average image of all motion-corrected images. Calcium fluorescence was extracted from all ROIs. Semi-automated data analysis was performed using custom Matlab (MathWorks) software. For each ROI, we corrected for potential neuropil contamination as previously described^84^. The ΔF/F (%) was calculated as the average change in fluorescence during the stimulus epoch relative to the 750 ms immediately prior to stimulus onset: ΔF/F (%)=((F_t_−F_0_)/F_0_)∗100. ROIs were included in additional analysis if they had a significant response (both p<0.05 Student’s two-tailed, paired t-test comparing activity during any stimulus and pre-stimulus epochs and had a mean ΔF/F equal to 5% or above for all trials with a particular frequency). Neurons were matched across sessions using the automated MATLAB algorithm ROIMatchPub (https://github.com/ransona/ROIMatchPub) with a matching criteria threshold of 0.4 (consistent with Seghal, et al., 2021) and manually verified (examples shown in **Extended Data** Fig. 5).

### Fiber photometry

Heterozygous ChAT-Cre mice were bilaterally injected with 1.0 µL of pAAV.Syn.Flex.GCaMP6s.WPRE.SV40 (diluted 1:2 in dPBS; Addgene: 100845-AAV5) in basal forebrain (AP: -0.5 mm, ML: -1.8 mm, DV: -4.5 mm from brain surface). Animals were head- posted and a 400 µm optical fiber (Thorlabs, Item# CFMC54L05) was implanted in the left hemisphere slightly above the injection site at -4.3 D-V. Experiments were performed three to four weeks after surgery. Photometry was performed using a custom-built rig (Falkner et al., 2016; Carcea et al., 2019). A 400 Hz sinusoidal blue light (40-45 µW) from a 470 nm LED (Thorlabs, Item# M470F1) connected to a LED driver (Thorlabs, Item# LEDD1B) was delivered via the optical fiber to excite GCaMP6s. We also used a control 330 Hz light (10 µW) from a 405 nm LED (Thorlabs, Item# M405FP1) connected to a second LED driver. Light travelled via 405 nm and 469 nm excitation filters via a dichroic mirror to the brain. Emitted light traveled back through the same optical fiber via dichroic mirror and 525 nm emission filter, passed through an adjustable zooming lens (Thorlabs, Item# SM1NR05) and was detected by a femtowatt silicon photoreceiver (Newport, Item# 2151). Recordings were performed using RX8 Multi-I/O Processor (Tucker-Davis Technologies). The envelopes of the signals were extracted in real time using Synapse software (Tucker-Davis Technologies). The analog readout was low-pass filtered at 10 Hz. Additional analysis was performed using custom MATLAB (Mathworks) software.

### Circuit tracing

For monosynaptic pseudotype tracing studies, 0.5 µL of a mix of pAAV-TREtight0mTag-BFP2- B19G (diluted 1:20 in dPBS; Addgene: 100799-AAV1) and pAAV-syn-FLEX-splitTVA-EGFP-tTA (diluted 1:200 in dPBS; Addgene: 100798-AAV1) was injected into basal forebrain (AP: -0.5 mm, ML: -1.8 mm, DV: -4.5 mm from brain surface) of ChAT-Cre mice. After one week, 0.25 µL of EnvA G-Deleted Rabies-mCherry (diluted 1:5 in dPBS; Addgene: 32636) was injected in the basal forebrain using the same coordinates. For mapping potential connectivity between basal forebrain, locus coeruleus and nucleus tractus solitarius, 0.75 µL of rgAAV-FLEX-tdTomato (diluted 1:3 in dPBS; Addgene number: 28306) was injected in TH-Cre mice in basal forebrain (AP: -0.5 mm, ML: -1.8 mm, DV: -4.5 mm from brain surface) using either a Hamilton syringe (5 µL) or Nanoject (Drummond Scientific; Part number: 3-000-207).

Animals were deeply anaesthetized with isoflurane and then transcardially perfused with phosphate buffered saline (1x PBS) followed by 4% paraformaldehyde (PFA) in PBS. Animals injected for circuit tracing studies were perfused 3-6 weeks after viral injection. Animals injected for optogenetics behavioral experiments were perfused after completion of behavioral training. After at least 12 hours in 4% PFA, brains were either transferred to PBS for sectioning with a vibratome or to a 30% sucrose-PBS solution for 24-48 hours to prepare for cryosectioning. For cryosectioning, brains were embedded in Tissue-Plus^TM^ O.C.T. Compound medium (Thermo Fisher Scientific, Item# 23-730) and sectioned using a cryostat (Leica). All sections were cut at 50 µm.

For animals injected with monosynaptic pseudotyped rabies, brain sections were washed with PBS (3x10 min at room temperature) and incubated for 2 hours at room temperature in blocking solution containing 5% normal goat serum (Millipore Sigma, Item # G6767) in 1% Triton X-100 (Millipore Sigma, Item #11332481001) dissolved in PBS. Brain slices containing basal forebrain were incubated in primary antibody (1:500 dilution of 3% normal goat serum in 1% Triton X100 dissolved in PBS of chicken anti-GFP IgY, Abcam catalog # ab13970 and rabbit anti-mCherry IgG, Abcam catalog # ab167453) for 24-48 hours at 4°C. Afterwards, slices were washed and incubated for 1–2 hours at room temperature in secondary antibody (1:500 dilution in PBS, goat anti-chicken IgY Alexa Fluor 488, Thermo Fisher Scientific cat. # A11039 and goat anti-Rabbit IgG (H+L) Cross-Absorbed Secondary Antibody, Alexa Fluor 555, Thermo Fisher Scientific Part number # A21428). Brain slices containing locus coeruleus and NTS were incubated in primary antibody (1:500 dilution of 3% normal goat serum in 1% Triton X100 dissolved in PBS, rabbit anti-mCherry IgG, Abcam catalog # ab167453) for 24-48 hours at 4°C. Slices were washed and incubated for 1–2 hours at room temperature in secondary antibody (1:500 dilution, goat anti-Rabbit IgG (H+L) Cross-Absorbed Secondary Antibody, Alexa Fluor 555, Thermo Fisher Scientific Part number # A21428). Finally, slides were washed and coverslipped using VECTASHIELD Antifade Mounting Medium with DAPI (Vector Labs Part number#: H- 1200-10). For all other animals, brain sections were washed with PBS and mounted using VECTASHIELD Antifade Mounting Medium with DAPI. Slides were imaged using a Carl Zeiss LSM 700 confocal microscope with four solid-state lasers (405/444, 488, 555, 639 nm) with appropriate filter sets and/or an Olympus AS-VSW whole slide scanner with an X-cite 120LED Boost High-Power LED Illumination system.

Individual brain sections were aligned with the Allen Brain Mouse Atlas^85^ using the QuickNII system^86^. The QuickNII system uses manual annotation and semi-automatic spatial registration to transform the reference atlas to match the anatomical landmarks in the corresponding experimental images. We then focused our quantification analysis to specific brain areas of interest – basal forebrain (AP: -0.5 mm), locus coeruleus (AP: -5.34) and solitary nucleus (AP: -6.36). Using the brain region outlines generated by QuickNII, we manually counted the number of cell bodies in each area. All labeled cells were marked in dots in shades of gray on the reference image for the corresponding anatomical plane.

### Optogenetic stimulation or inhibition of basal forebrain cholinergic neurons

To stimulate auditory cortical projecting cholinergic neurons, 1.0 µL of AAVrg.EF1a. doublefloxed.hChR2(H134R).EYFP.WPRE-HGHpA (diluted 1:2 with dPBS; Addgene number: 20298) was injected into auditory cortex as described above in ChAT-Cre animals. For optogenetic inhibition, 0.50 µL of AAV5.FLEX.ArchT.tdTomato (diluted 1:2 with dPBS; Addgene number: 28305) was injected in basal forebrain of ChAT-Cre animals using the coordinates described above. In both cases, optic fibers were implanted 50 µm above basal forebrain and the animal was head-posted. Optic fibers were custom made of glass fibers (200 μm core; Thorlabs FT200UMT) fitted with zirconia LC connectors (Precision Fiber Products MM-FER2007C-2300), secured using glue (Krazy Glue). All fibers used had at least an 80% efficiency prior to implantation.

For optogenetic stimulation during behavior, cholinergic basal forebrain neurons were bilaterally stimulated in the same way as VNS pairing (500 ms centered around the tone at 30 Hz; two blocks of 50 stimulation trials surrounded by unstimulated trials). For optogenetic inactivation experiments, cholinergic basal forebrain neurons were bilaterally inhibited using a 500 ms pulse, presented in conjunction with VNS pairing. Light (either blue or yellow) was delivered using a driver (Thorlabs LEDD1B), fiber coupled LED (Thorlabs M470F3 or M565F3), a patch cable (Thorlabs M129L01), and a bifurcated fiber bundle (Thorlabs BFYL2LF01) connected to the fibers using a mating sleeve (Thorlabs ADAL1). Laser power was calibrated across days and between 1-3 mW for all optogenetics experiments during behavior.

## Author Contributions

K.A.M., J.K.S and R.C.F. designed the experiments. K.A.M, E.S.P, and J.K.S. performed the experiments. K.A.M. performed analysis. E.S.P., S.S.F., and S.O.V. provided technical assistance. N.Z.T., E.S.P., D.A.M., M.J.M., and R.C.F. designed and validated the VNS cuff electrode. K.A.M and R.C.F. wrote the manuscript.

## Acknowledgments

We thank A. Agha, C.L. Ebbesen, E. Glennon, E. Kay-Rivest, R.A. Kolaric, K.I. Nagel, D.H. Sanes, D.M. Schneider, M.A. Svirsky, J.T. Roland, and S. Valtcheva for comments, discussions, and/or technical assistance. We thank the GENIE Program, Janelia Research Campus, and the Howard Hughes Medical Institute for provision of GCaMP6s and ScanImage. This work was funded by grants from the Defense Advanced Research Projects Agency (N66001-17-2-4010 to R.C.F., D.A.M., and M.J.M.), the National Institute on Deafness and Other Communication Disorders (DC012557 to R.C.F.), and the National Science Foundation (to K.A.M. and J.K.S.).

---

**Extended Data Figure 1. Performance is stable and does not significantly improve after seven days in stage 3. a**, Lick rates relative to tone offset from one session in six fully trained example animals (shades of gray represent individual animals). **b**, Lick rates in the second after tone offset (2.6±0.84 licks/s, mean±s.d., N=6) are significantly higher than lick rates 2-3 seconds after tone offset (0.2±0.3 licks/s, p=0.0002, Student’s two-tailed paired t-test, N=6). **c**, Change in performance across all frequencies throughout all days in stage 3 relative to the performance on the first three days of stage 3 (N=38). **d**, Correlation between total days animals spend in stage three and peak performance in stage 3 (Pearson’s R=0.31, p=0.06, N=38). **e**, Mean performance significantly improved across days (Days 0-2: 64.7±1.0%; Days 6-8: 70.0±1.2%; Days max±1: 72.6±1.3%, mean±s.d.; p<0.00002 one-way ANOVA with Tukey’s multiple comparisons correction, N=38 animals). **f**, Across-animal performance is variable on day 7 of stage three (69.0±8.0%, mean±s.d., N=38 animals). **g**, Percent reported center at each frequency relative to center stimulus on days 0- 2, days 6-8 and behavior on day of maximum performance in stage 3. **h**, Error rate at frequencies 0 (green; day 1-3: 18.7±2.3%, day 7-9: 18.3±2.0%, Max day ± 1 day: 15.0±1.8%, mean±s.e.m.; p=0.36, one-way ANOVA with Tukey’s multiple comparisons correction), ±0.25 (dark gray; day 1-3: 79.2±2.4%, day 7-9: 71.0± 2.7%, Max±1: 73.0±2.5%, mean±s.e.m.; p=0.07, one-way ANOVA with Tukey’s multiple comparisons correction) and ±0.5 (medium gray; day 1-3: 69.8±2.9 %, day 7-9: 56.2±2.7%, Max±1: 53.6±2.9%, mean±s.e.m.; p=0.0002, one-way ANOVA with Tukey’s multiple comparisons correction) from center over days 0-2 (open circles, white), 6-8 (open circles, gray), and max (± 1 day, closed circles) in stage 3.

**Extended Data Figure 2. Additional training did not continue to improve performance. a**, Percent reported left for all days in stage 3 for animals trained for 21-48 days in stage 3 (36.3±12.4 days in stage 3, N=6). ND (gray) represents days with no data collection. **b**, Schematic for reward size modulation experiments performed in days following performance shown in **a**. After animals were fully trained and had stable performance, animals received rewards of double the volume for all frequencies for 12-13 additional days. **c**, Performance over all frequencies over days of altered reward size. **d**, Average percent reported center for the five days prior to increased reward size and last five days on increased reward size. **e**, Performance across the two reward sizes (smaller reward, more trials: open circle, larger reward, fewer trials: closed circle) at 0 (small reward: 78.9±4.1% correct, for large reward: 83.5±3.3%, mean±s.e.m; p=0.15, Student’s two-tailed paired t-test), ±0.25 (small reward: 36.9±6.8% correct, large reward: 35.4±4.3%, mean± s.e.m.; p=0.67, Student’s two-tailed paired t-test), and ±0.5 (small reward: 64.2±7.0% correct, large reward: 71.4±6.3%, mean± s.e.m.; p=0.08, Student’s two-tailed paired t-test).

**Extended Data Figure 3. VNS transiently alters heart rate in viable cuffs. a,** Impedance over days post-implantation from a representative animal (top) and all 7 wild-type mice used for VNS pairing behavioral experiments. Gray, individual animals; black, mean±s.d. every 10 days (N=7 mice). **b**, Cuff impedance is stable over time. Impedance reading on day 0 (day of cuff implantation) and the first measurement 30+ days after implantation (30- 54 days) (Day 0 mean: 2.7±0.6 kΩ, day 30-54 mean: 2.3±1.6 kΩ, mean±s.d., N=7 mice, p=0.56, Student’s two-tailed paired t-test). **c**, Description of vitals recordings. Sessions with and without VNS application were alternated. Between 10 and 20 VNS bouts occurred in each VNS session and lasted 500 ms at 0.1 Hz. Stimulation intensity was consistent within a session but was systematically changed throughout the day (ranging from 0.2 to 1.4 mA). The following data is from sessions with VNS (0.8-1.0 mA) compared to the baseline sessions immediately prior or following. **d**, Distribution of heart rate was not significantly different during VNS or baseline sessions in two animals with cuff impedances >100MΩ (Sham 1, cuff impedance: 100MΩ, p=0.66, Mann Whitney U two-sided test; Sham 2, cuff impedance: 120MΩ, p=0.48, Mann Whitney U two-sided test). **e**, Raw heart rate from example animal with viable cuff. VNS was applied at 0.1 Hz and lasted 500 ms. **f**, Distribution of heart rates in VNS and baseline sessions for ten animals with potentially viable cuffs (VNS significantly reduced heart rate in 7/10 animals; Animal 1, cuff impedance: 0.2 kΩ, p<10^-5^, Mann Whitney U two-sided test; Animal 2, cuff impedance: 3.4 kΩ, p<10^-5^; Animal 3, cuff impedance: 3.0 kΩ, p<10^-5^; Animal 4, cuff impedance: 0.1 kΩ, p=0.03; Animal 5, cuff impedance: 2.8 kΩ, p<10^-5^; Animal 6, cuff impedance: 3.9 kΩ, p<10^-5^; Animal 7, cuff impedance: 3.9 kΩ, p<10^-5^; Sham 3, cuff impedance: 2.1 kΩ, p>0.05; Sham 4, cuff impedance: 1.9 kΩ, p>0.05; Sham 5, cuff impedance: 3.6 kΩ, p>0.05).

**Extended Data Figure 4. Experimental and control animals have consistent behavior prior to VNS pairing. a**, Distribution of days in stage one for sham control (gray, 13.9±6.0 days, mean±s.d., N=10) and experimental animals (black, 10.4±4.4 days, mean±s.d., N=7, p=0.16, Mann Whitney U two-sided test). **b**, Distribution of days spent in stage 3 prior to VNS pairing for sham control (gray, 17.5±6.8 days, mean±s.d., N=10) and experimental animals (black, 21.1±10.6 days, mean±s.d., N=7, p=0.40, Mann Whitney U two-sided test). **c**, Distribution of response rate in stage 3 for sham control (gray, 96.1±5.0%, mean±s.d., N=10) and experimental animals (black, 91.3±7.5%, mean±s.d., N=7, p=0.14, Mann Whitney U two-sided test). **d**, Distribution of peak performance in stage 3 for sham control (gray, 73.4±6.1%, mean±s.d., N=10) and experimental animals (black, 68.7±10.0%, mean±s.d., N=7, p=0.36, Mann Whitney U two-sided test). **e**, Percent reported center for sham (dotted lines, N=10) and experimental animals (solid lines, N=7, p=0.20, two-way ANOVA with Tukey’s multiple comparisons correction). **f**, Performance is not affected by VNS cuff implantation in experimental (-1.8±1.7%, mean±s.e.m., N=7, p=0.38, Student’s two-tailed paired t-test) or sham animal (0.4±1.0%, mean±s.e.m., N=10, p=0.68, Student’s two-tailed paired t-test; difference between sham and experimental: p=0.28, Student’s two-tailed unpaired t-test).

**Extended Data Figure 5. Impact of VNS pairing on experimental and control animals. a**, Change in response rate for the three days prior to VNS onset and on days 18-20 of VNS pairing for sham (gray, -7.8±2.4% response rate, mean±s.e.m., N=10) and experimental animals (black, 4.5±2.1% response rate, mean±s.e.m., N=7, p=0.002, Student’s two-tailed unpaired t-test). **b**, Response rate across frequencies for the three days prior to VNS onset (black) and on days 18-20 of VNS pairing (blue) for experimental animals (p=0.32 across frequencies, p<0.001 across days, ANOVA with Tukey’s multiple comparisons correction). **c**, Percent correct on final days of VNS pairing (Days 18-20) across all frequencies in experimental animals (blue, closed circles, without: 80.4±2.4%, mean±s.e.m., with: 77.1±3.1%, N=7) in comparison to the behavior three days prior to VNS (black, closed circles, ‘Day -2-0’, without: 71.6±3.5%, with: 65.4±3.8%) with (p=0.008, Student’s paired t-test) and without no response trials (without: p=0.02, Student’s paired t-test). **d**, Change in response rate for the three days prior to VNS onset and days 18-20 of VNS pairing across three groups of intensities used for control animals (p=0.4, one-way ANOVA with Tukey’s multiple comparisons correction, N=10). **e**, Percent of trials reported as center for all animals used in VNS pairing experiments for the three days prior to VNS onset (black) and on days 18-20 of VNS pairing (blue) used in Figure 2 and 3 (Control: N=10, dotted lines; Experimental: N=7, solid lines).

**Extended Data Figure 6. Tracking neurons during two-photon imaging. a**, Example ROIs in an example animal during unpaired passive tone presentation on days 0, 3, 6, 9 and 11 days of VNS pairing. Average Pearson’s R correlation for 20x20 pixel area containing ROIs and surrounding area across all days. **b**, Average Pearson’s R correlation for 20x20 pixel area containing ROIs and surrounding area across all days of passive tone presentation (“Imaging days”) for all VNS pairing animals (N=7; Average R (animal 1): 0.59±0.15, mean±s.d.; Average R (animal 2): 0.67±0.12; Average R (animal 3): 0.69±0.08; Average R (animal 4): 0.52±0.18; Average R (animal 5): 0.69±0.13; Average R (animal 6): 0.61±0.15; Average R (animal 7): 0.57±0.19).

**Extended Data Figure 7. Activation of basal forebrain cholinergic neurons by VNS. a**, Correlation between baseline activity of cholinergic cell bodies in 1.5s prior to VNS onset and average activity in 1.5s following VNS onset (Pearson’s R=-0.30, p=0.02, N=3 mice, n=60 trials). **b**, Cholinergic cell body responses prior to or during VNS separated by baseline responses (higher than zero or lower than or equal to zero). Responses in 500ms after VNS were significantly higher in trials with low baseline (during VNS: 0.47±0.16% ΔF/F, p=0.006, Student’s two-tailed t-test) but not during trials with high baseline (during VNS: -0.21±0.19% ΔF/F, p=0.26, Student’s two-tailed t-test). **c**, Example basal forebrain and NTS section with alignment to corresponding output from QuickNII system of Allen Brain Atlas section with either basal forebrain or NTS circled in white. **d**, Schematic for mapping inputs to cholinergic basal forebrain neurons using retrograde, Cre-dependent, pseudotyped monosynaptic rabies. Location of mCherry labeled neurons in section containing LC labeled by dots in shades of gray. The region of interest is highlighted in red. Each animal is represented in a distinct shade of gray (N=5). **e**, Schematic of injection of retrograde Cre-dependent tdTomato into basal forebrain of TH-Cre mice. Location of tdTomato labeled neurons in section containing NTS labeled by dots in shades of gray. The region of interest is highlighted in red. Each animal is represented in a distinct shade of gray (N=4). **f**, Schematic of proposed anatomical connections between NTS, locus coeruleus and basal forebrain. **g**, Correlation between baseline cholinergic axon activity in 1s prior to VNS onset and average activity in the 1s following VNS onset (Pearson’s R=0.04, p=0.71).

**Extended Data Fig. 8. Impact of stimulating auditory cortical projecting cholinergic neurons on individual animal performance. a**, Verification of fiber placement over basal forebrain in one example animal. Scale bar is 1200 μm. **b**, Distribution of peak performance prior to optogenetic pairing onset of control (open circles, 77.1±2.5% correct, mean±s.e.m.) and experimental animals (‘ChR2’, closed circles, 72.1±2.8% correct, mean ± s.e.m., p=0.15, Mann Whitney U two-sided test). **c**, Percent of trials reported center for control (dotted lines, mean in blue, individual animals in gray, N=7) and experimental animals (solid lines, mean in blue, individual animals in gray, N=8). **d**, Difference in response rate in the three days prior to optogenetic or sham pairing and on the last three days of optogenetic or sham pairing in control (open circles, -0.28±7.5% change in response rate, mean±s.e.m, N=7) or experimental animals (closed circles, -1.7±1.8% change in response rate, mean±s.e.m., N=8, p=0.86, Student’s two-tailed unpaired t-test). **e**, Percent of trials reported as center for 15 animals in Figure 6 for the three days prior to optogenetic or sham pairing onset (black) and on the last three days with optogenetic or sham pairing (blue). Experimental animals are represented with solid lines and sham animals are represented with dotted lines.

**Extended Data Figure 9. Impact of inhibiting cholinergic neurons in basal forebrain during VNS on individual animal performance. a**, Verification of fiber of fiber placement over basal forebrain in one example animal. Scale bar is 1200 μm. **e**, Percent of trials reported as center for 6 animals in Fig. 7 for the three days prior to VNS onset (black) and days 18-20 of VNS stimulation (orange). **b,** Description of vitals recordings. Sessions with and without VNS application were alternated. Between 10 and 20 VNS bouts occurred in each VNS session and lasted 500 ms at 0.1 Hz. Stimulation intensity was consistent within a session but was systematically changed throughout the day (ranging from 0.2 to 1.4 mA). The following data is from sessions with VNS (0.8-1.0 mA) compared to the baseline sessions immediately prior or following. **c**, Example baseline and VNS session from example animal. **d**, Distribution of heart rates in VNS (blue) and baseline (gray) sessions for six animals with potentially viable cuffs (p<0.001, Mann Whitney U two-sided test). **e**, Difference in response rate in the three days prior to VNS pairing and on day 18-20 of VNS pairing with inhibition of cholinergic basal forebrain neurons (1.4±3.3% change in response rate, mean±s.e.m, N=6). **f**, Percent of trials reported as center for 6 animals in Figure 7 for the three days prior to optoinhibition during VNS pairing onset (black) and on the last three days with optoinhibition during VNS pairing (orange).

## Notes

### Competing Interest Statement

The authors have declared no competing interest.

### Summary of Updates

First round of reviewer comments

