## Extended Data Figures for "Vagus nerve stimulation recruits the central cholinergic system to enhance perceptual learning"

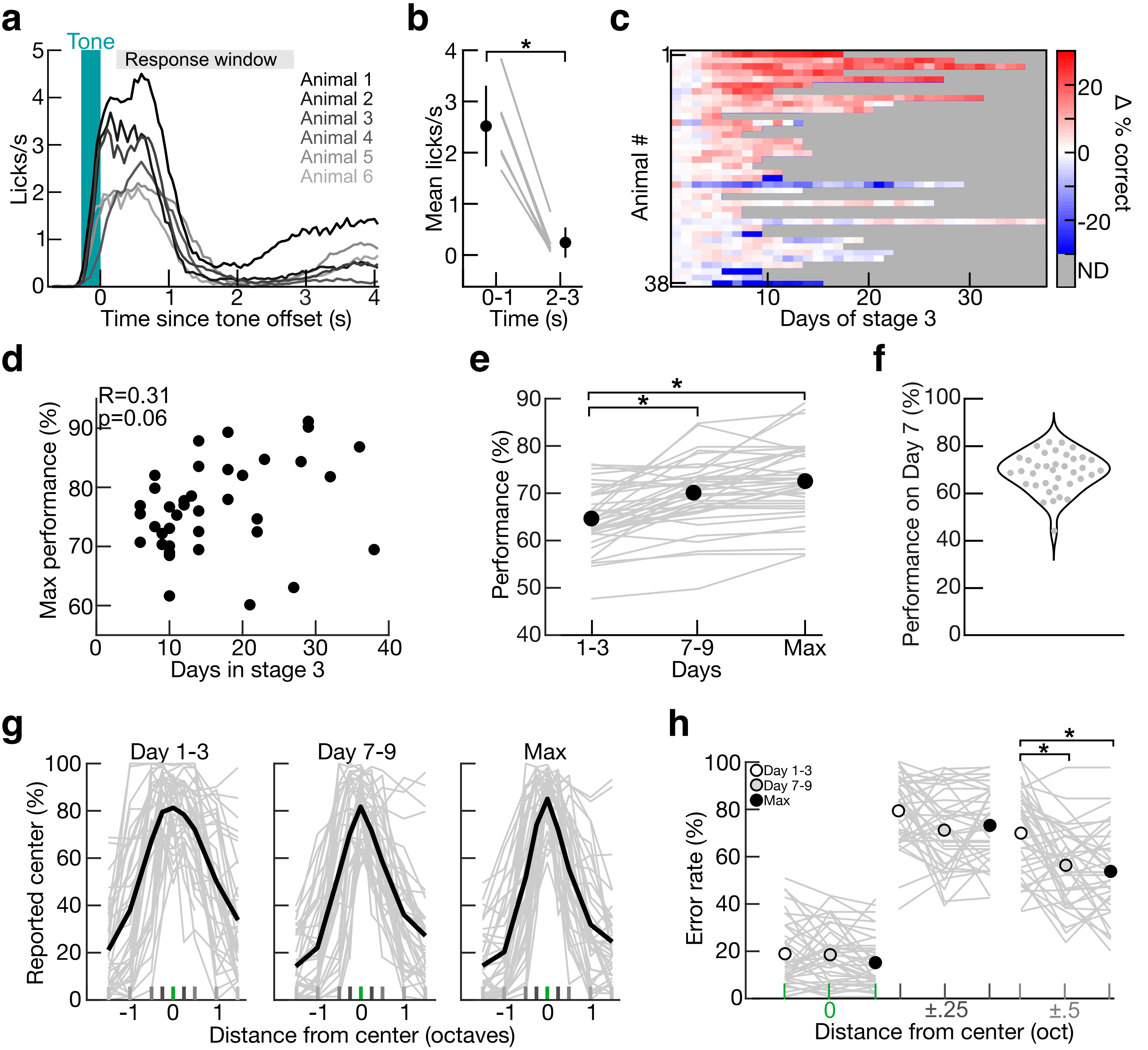


Extended Data Figure 1

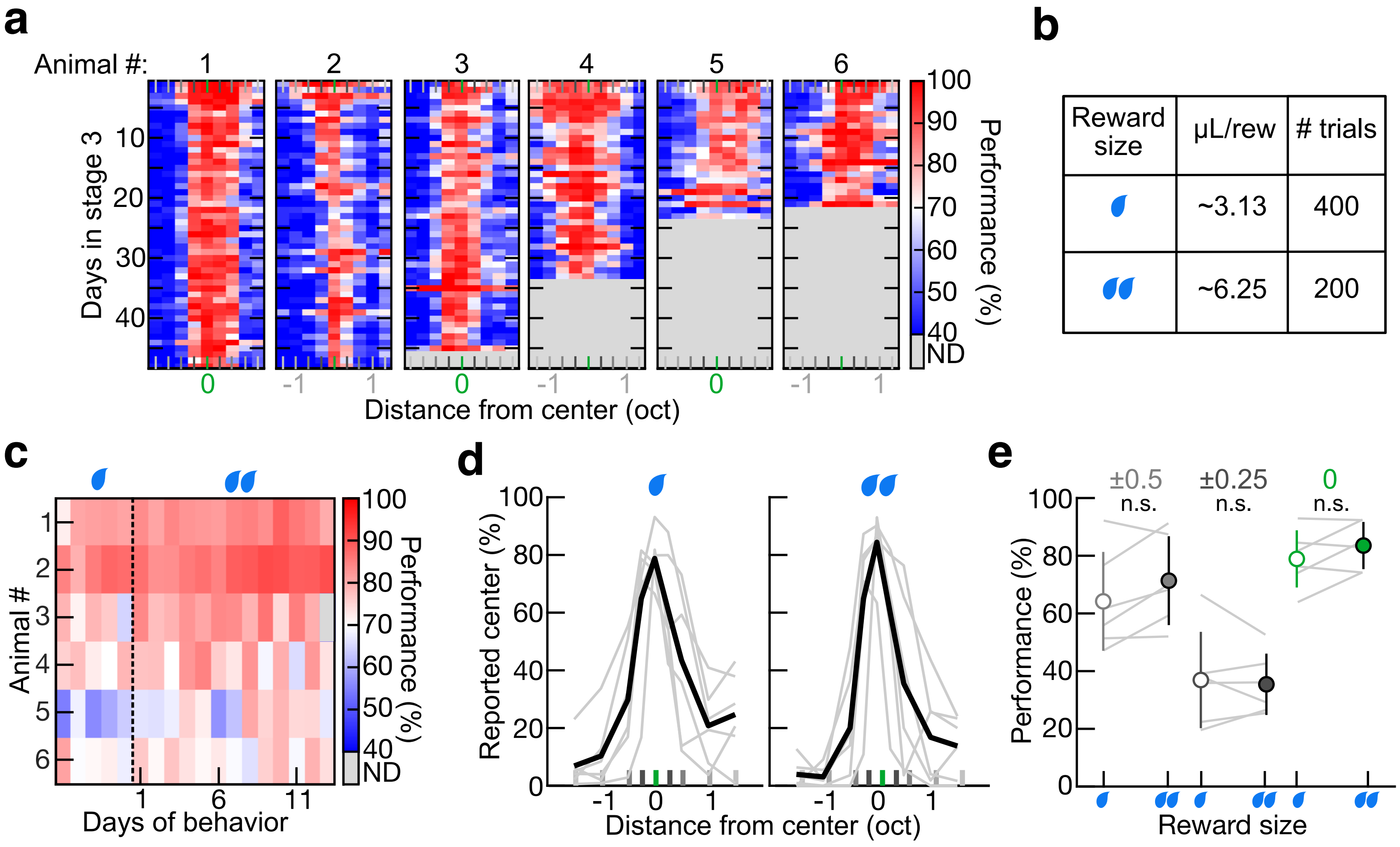


Extended Data Figure 2

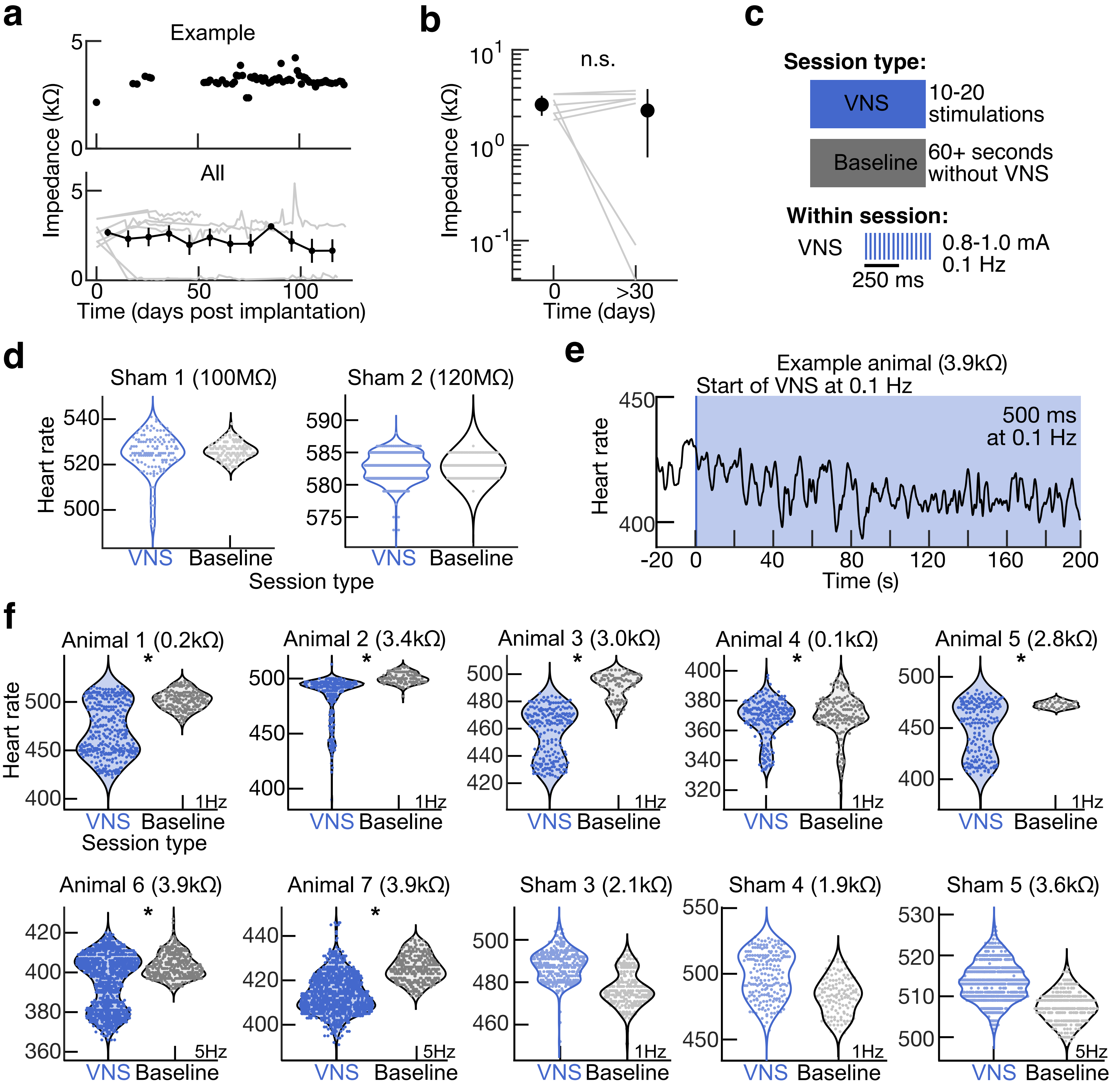


Extended Data Figure 3

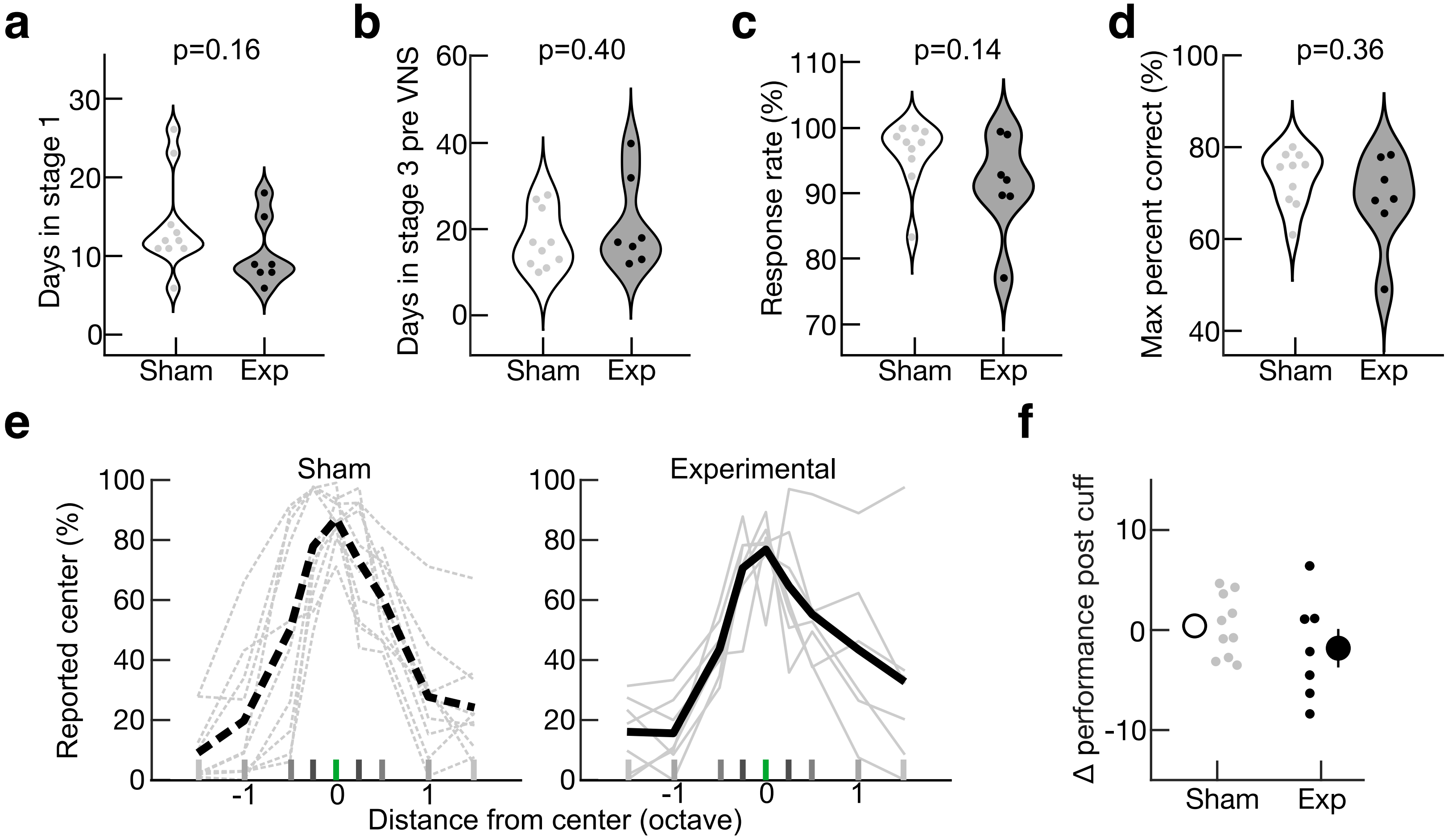


Extended Data Figure 4

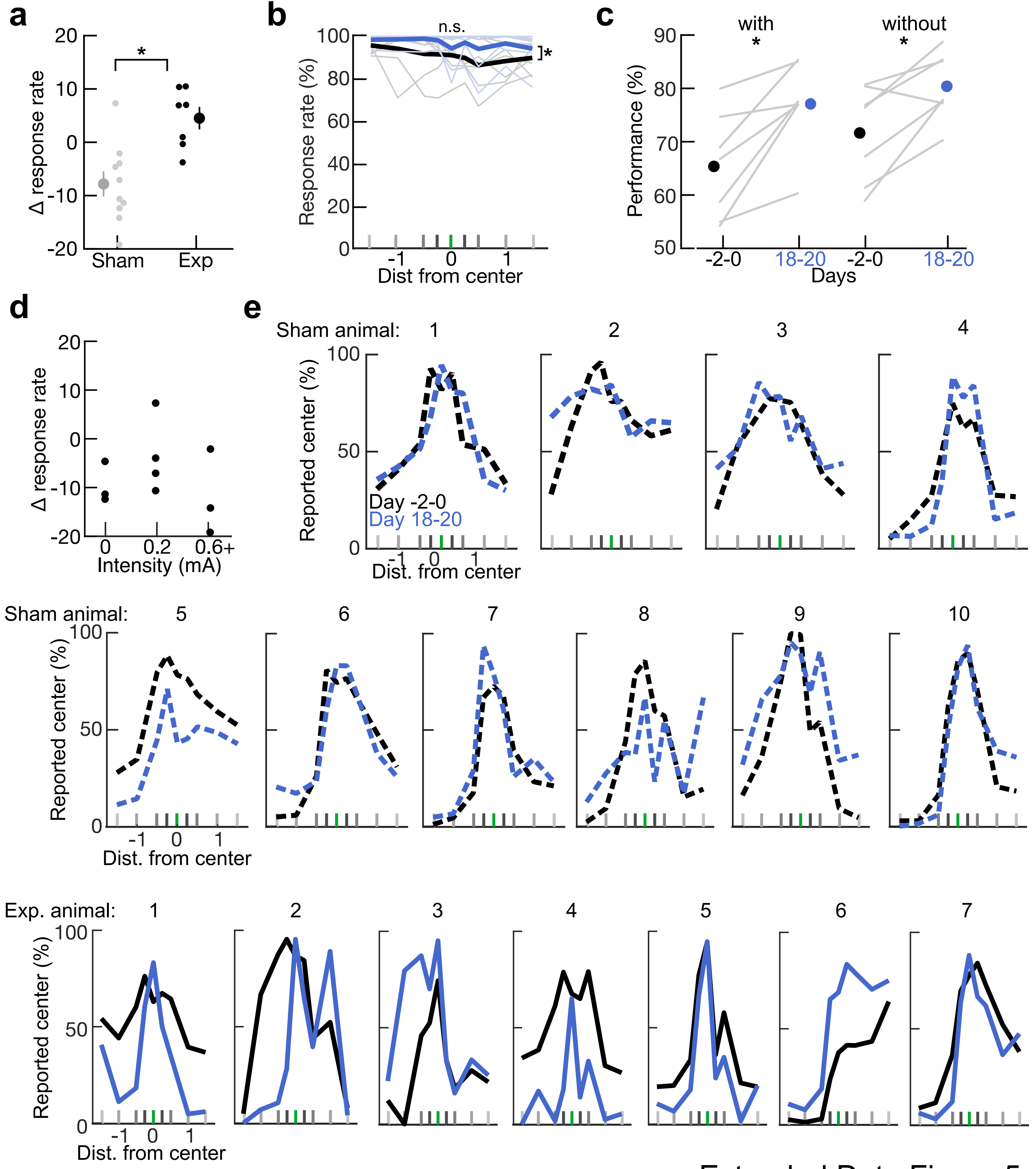


Extended Data Figure 5

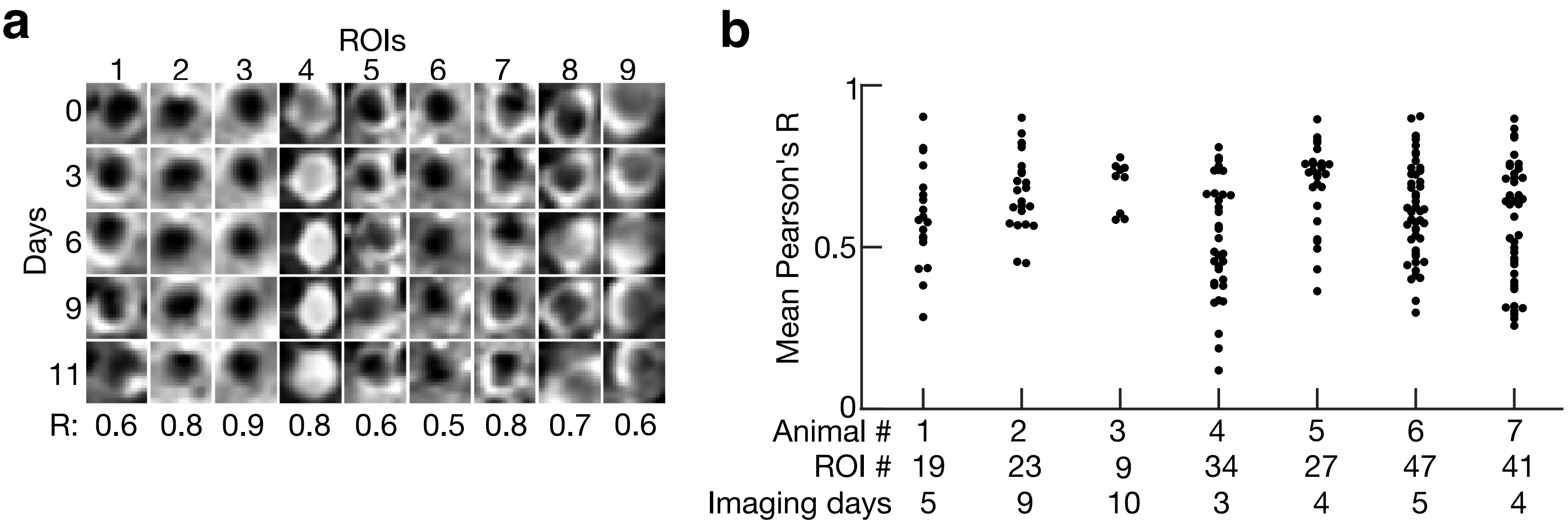


Extended Data Figure 6

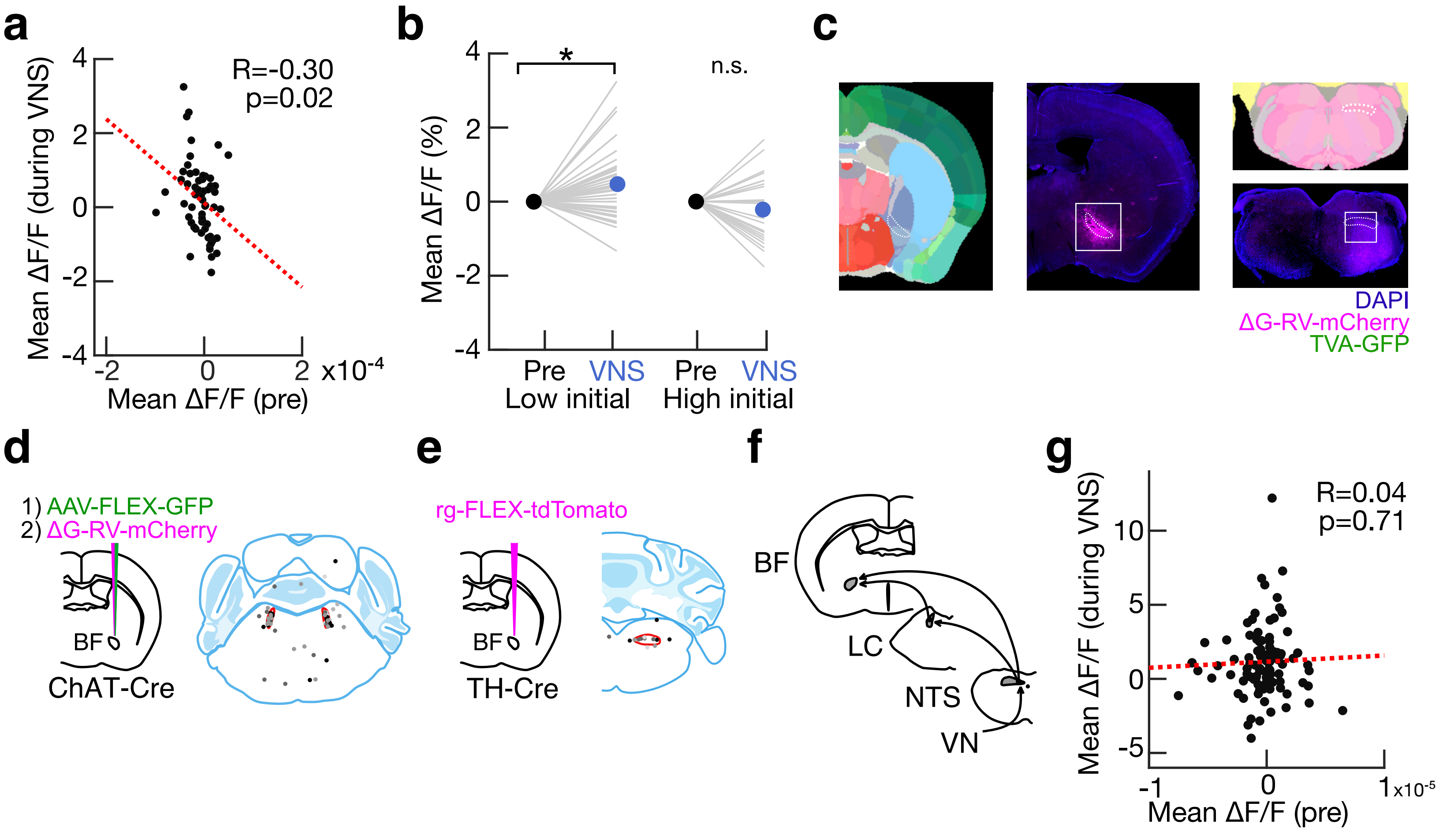


Extended Data Figure 7

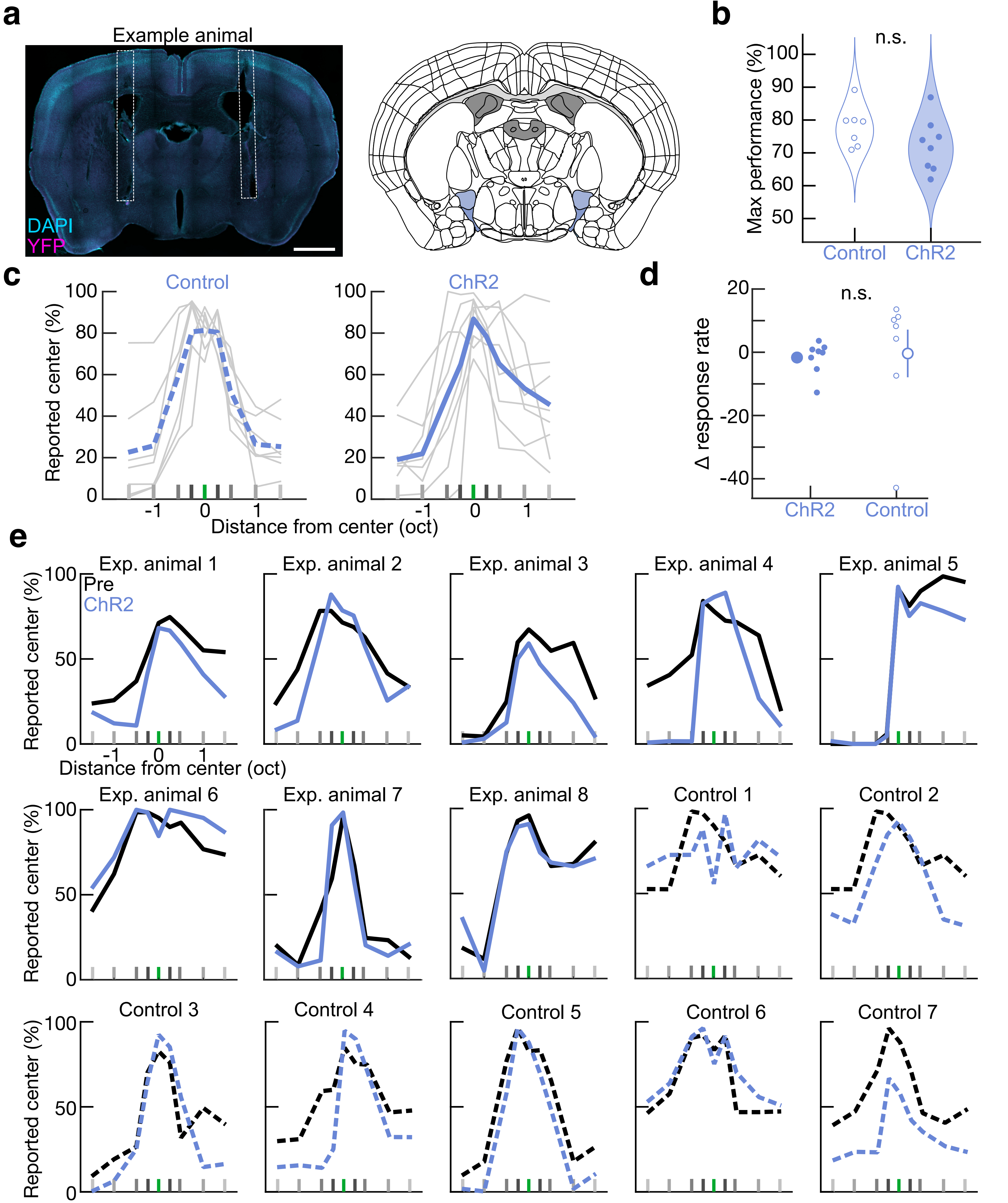


Extended Data Figure 8

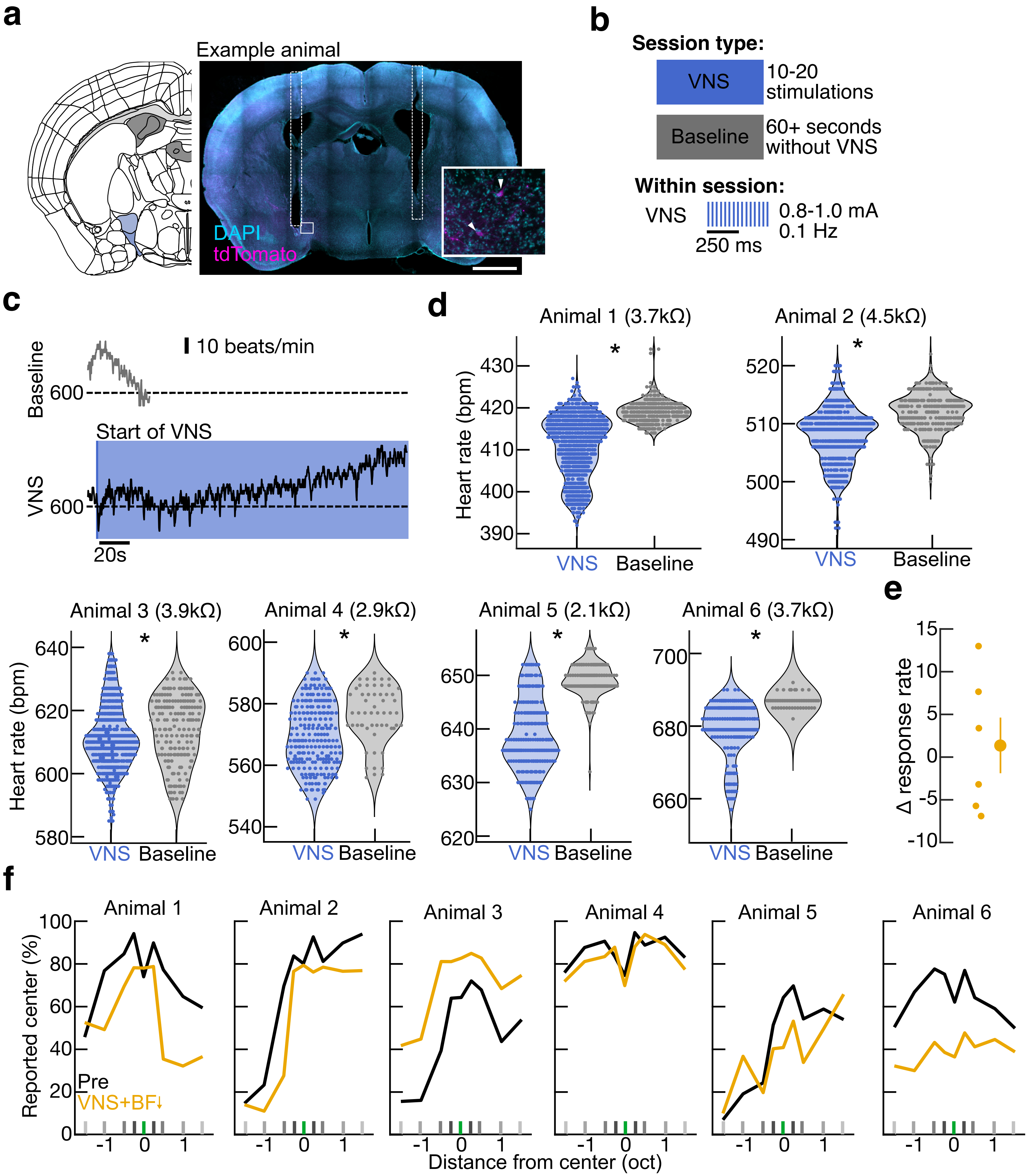


Extended Data Figure 9
